# The repurposed drugs suramin and quinacrine inhibit cooperatively *in vitro* SARS-CoV-2 3CL^pro^

**DOI:** 10.1101/2020.11.11.378018

**Authors:** Raphael J. Eberle, Danilo S. Olivier, Marcos S. Amaral, Dieter Willbold, Raghuvir K. Arni, Monika A. Coronado

## Abstract

Since the first report of a new pneumonia disease in December 2019 (Wuhan, China) up to now WHO reported more than 50 million confirmed cases and more than one million losses, globally. The causative agent of COVID-19 (SARS-CoV-2) has spread worldwide resulting in a pandemic of unprecedented magnitude. To date, no clinically safe drug or vaccine is available and the development of molecules to combat SARS-CoV-2 infections is imminent. A well-known strategy to identify molecules with inhibitory potential against SARS-CoV-2 proteins is the repurposing of clinically developed drugs, e.g., anti-parasitic drugs. The results described in this study demonstrate the inhibitory potential of quinacrine and suramin against SARS-CoV-2 main protease (3CL^pro^). Quinacrine and suramin molecules present a competitive and non-competitive mode of inhibition, respectively, with IC_50_ and K_D_ values in low μM range. Using docking and molecular dynamics simulations we identified a possible binding mode and the amino acids involved in these interactions. Our results suggested that suramin in combination with quinacrine showed promising synergistic efficacy to inhibit SARS-CoV-2 3CL^pro^. The identification of effective, synergistic drug combinations could lead to the design of better treatments for the COVID-19 disease. Drug repositioning offers hope to the SARS-CoV-2 control.

## 1. Introduction

December 2019, local health authorities reported an increasing number of pneumonia cases spreading rapidly across the city of Wuhan, in the Hubei province in China. The causative agent of this disease was identified as SARS-coronavirus-2 (SARS-CoV-2) (Lu et al., 2020) and although most cases are asymptomatic or only evidence mild symptoms with good recovery rates; a small percentage of the infected patients develop more severe manifestations, such as severe pneumonia, respiratory failure, multiple organ failures and multiple cases resulting in death (Huang et al., 2020). SARS-CoV-2 has spread worldwide, leading to a coronavirus pandemic of unprecedented magnitude and the World Health Organization (WHO) declared the coronavirus disease 2019 (COVID-19) as an international public health emergency (Lai et al., 2020). Globally, more than 50 million confirmed cases resulted in over 1.2 million victims as reported by the WHO on 11th of November 2020 (World Health Organization, 2020). This staggering number is a major challenge to precarious healthcare systems, especially, in developing countries. Under these circumstances, the identification and development of safe and effective SARS-CoV-2 drugs is of high priority.

Coronoviridae forms the largest family of positive-sense single-stranded RNA viruses and is classified into four genera (α, β, γ, and δ) (Nga et al., 2011). SARS-CoV, Middle East respiratory syndrome coronavirus (MERS-CoV), and SARS-CoV-2 are β-coronaviruses (de Wit et al., 2020). Analysis of the SARS-CoV-2 genome sequences revealed a higher identity to bat SARS-like coronavirus (89.1% nucleotide similarity) (Wu et al., 2020).

The genomic RNA of coronaviruses is ~30,000 nucleotides in length and contains at least six open reading frames (ORFs) (Hussain et al., 2005; Chen et al., 2020) The first ORF (ORF 1a/b), about two-thirds of the genome length, directly translates two polyproteins, pp1a and pp1ab, so named because there is an a-1 frameshift between ORF1a and ORF1b. These polyproteins are processed by a main protease, M^pro^ (also known as 3C-like protease - 3CL^pro^) and, by viral papain-like proteases into 16 nonstructural proteins (NSPs). These NSPs are involved in the generation of subgenomic RNAs that encode four main structural proteins (envelope (E), membrane (M), spike (S), and nucleocapsid (N) proteins) and other accessory proteins (Ramajayam et al., 2011; Ren et al., 2013). Therefore, these proteases, especially 3CL^pro^, play vital roles in the replication process of coronaviruses.

3CL^pro^ is a cysteine protease with a three-domain (domains I to III) organization and with a chymotrypsin like fold (Anand et al., 2002). Active 3CL^pro^ consists of a homodimer containing two protomers and the coronavirus 3CL^pro^ features a noncanonical Cys-His dyad located in the cleft between domains I and II (Anand et al., 2002; Anand et al., 2003; Yang et al., 2003).

The functional importance of 3CL^pro^ in the viral life cycle combined with the absence of closely related homologues in humans, indicating that this protease is an attractive target for the development of antiviral drugs (Pillaiyar et al., 2016). So far, there are no efficacious drugs and vaccines available to combat SARS-CoV-2 infections and the bureaucratic process to approve new drugs is a very time consuming and costly process. Thus, repurposing of existing drug molecules could be a rapid alternative to combat the virus infection. The well described anti-parasitic drugs like chloroquine, quinacrine and suramin revealed anti-viral effects against SARS-Cov-2 in cell cultures (Liu et al., 2020; da Silva et al., 2020; Duan et al., 2020; Roldan et al., 2020; Touret and de Lamballerie, 2020). These three compounds have been used as antiprotozoal drugs, being that chloroquine and quinacrine are used against malaria (Hoekenga, 1955; Joy, 1999) and suramin against sleeping sickness, which is caused by trypanosomes (Steverding, 2010).

In the present study, we tested the inhibitory potentials of chloroquine, quinacrine and suramin against SARS-CoV-2 3CL^pro^. Quinacrine and suramin presented IC_50_ values < 10 μM and K_D_ values in a low μM range; however, chloroquine did not affect the SARS-CoV-2 3CL^pro^ activity. Further experiments identified quinacrine and suramin as competitive and non-competitive inhibitor, respectively. A 1:1 combination of quinacrine and suramin decreased the IC_50_ value to < 500 nM. Our results indicate that quinacrine and suramin are interesting drug candidates to combat SARS-Cov-2 infections and molecular docking and molecular dynamics simulation inferred the amino acid residues responsible for the interaction between the 3C-like protease and the molecules.

## 2. Materials and Methods

### 2.1 Cloning, expression and purification of SARS-CoV-2 3CL^pro^

The codon optimized cDNA encoding SARS-CoV-2 3CL^pro^ (Uniprot entry: P0DTD1) was synthesized and implemented in the ampicillin resistant vector pGEX-6P-3 (BioCat GmbH, Heidelberg, Germany). The construct contains an N-terminal GST-tag and a PreScission protease cleavage site (LEFLFQGP).

SARS-CoV-2 3CL^pro^-pGEX-6P-3 vectors were transformed into *E. coli* Lemo21 (DE3) (New England BioLabs, USA) competent cells and has grown overnight at 37 °C in LB-medium. This pre-culture was added to fresh LB-medium (Ampicillin and Chloramphenicol) and grew at 37 °C until the cells reached an OD600 of 0.6. Gene expression was induced with final concentration of 0.5 mM IPTG (1 mM Rhamnose was added) and incubated for 3h, at 37 °C and 120 rpm. Subsequently, the culture was harvested by centrifugation (4,000 rpm) at 5 °C for 20 min (Sorvall RC-5B Plus Superspeed Centrifuge, Thermo Fisher Scientific, USA; GSA rotor). The supernatant was discarded and the cells containing the recombinant SARS-CoV-2 3CL^pro^_GST was resuspended in 50 mM Tris-HCl pH 8.0, 200 mM NaCl (lysis buffer) and stored at −20 °C for subsequent purification.

For purification, the cell-suspension was incubated on ice for 1 h with addition of lysozyme, subsequently it was lysed by sonication in four pulses of 30 s each with amplitude of 30% interspersed by intervals of 10 s. The crude cell extract obtained was centrifuged (7,000 rpm for 90 min at 6 °C). The supernatant containing SARS-CoV-2 3CL^pro^_GST was loaded onto a GSH-Sepharose matrix which was previously extensively washed with the lysis buffer and was extensively washed with the same buffer. The protein was eluted with the same buffer plus addition of 10 mM GSH. The eluted fractions were concentrated and dialyzed against PreScission protease cleavage buffer (50 mM Tris pH 7.0, 200 mM NaCl, 1 mM DTT and 1 mM EDTA). PreScission protease was used to cleave the GST-tag from the SARS-CoV-2 3CL^pro^_GST fused protein. For 100 μg target protein concentration, 10 μg PreScission protease was added and the sample was incubated for 36 h at 4 °C. Separation of the target protein, the GST-tag and the PreScission protease was achieved using GSH-Sepharose. Further, to remove aggregated fraction, size exclusion chromatography was used (Superdex 200 10/300 GL GE Healthcare, USA), the column was equilibrated with 20 mM Tris-HCL pH 8.0, 150 mM NaCl. Sample purity after each purification step was assessed by 15% SDS-PAGE gels. The corresponding protein fraction was concentrated up to 2 mg/ml and stored at −20 °C.

### 2.2 Activity assay of SARS-CoV-2 3CL^pro^

SARS-CoV-2 3CL^pro^ activity assay was performed as described earlier using a fluorogenic substrate DABCYL-KTSAVLQ↓SGFRKME-EDANS (Bachem, Switzerland) in a buffer containing 20 mM Tris pH 7.2, 200 mM NaCl, 1 mM EDTA and 1 mM TCEP (Zhang et al., 2020a; Zhang et al., 2020b; Ma et al., 2020). The reaction mixture was pipetted in a Corning 96-Well plate (Sigma Aldrich) consisting of 0.5 μM protein and the assay was initiated with the addition of the substrate at a final concentration of 50 μM. The fluorescence intensities were measured at 60 s intervals over 30 minutes using an Infinite 200 PRO plate reader (Tecan, Männedorf, Switzerland). The temperature was set to 37 °C. The excitation and emission wavelengths were 360 nm and 460 nm, respectively. For K_M_ and V_max_ measurements, the procedure was followed as described previously (Ma et al., 2020). A substrate concentration from 0 to 200 μM was applied. The initial velocity of the proteolytic activity was calculated by linear regression for the first 15 minutes of the kinetic progress curves. The initial velocity was plotted against the substrate concentration with the classic Michaelis-Menten equation using GraphPad Prism^5^ software and Kcat was obtained using the equation (1):

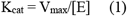

while V_max_ is the experimental determined maximal velocity and [E] is the enzyme concentration in the experiment (Berg et al., 2002). All measurements were performed in triplicate and data are presented as mean ± SD.

### 2.3 Inhibition assay of SARS-CoV-2 3CL^pro^

Inhibition of SARS-CoV-2 3CL^pro^ activity by chloroquine, quinacrine and suramin was investigated using the activity assay described above. 10 μM of the compounds was used for a preliminary screening test. For the final inhibition assays 0.5 μM of the protein was incubated with 0-100 μM suramin and 0-150 μM quinacrine, chloroquine did not show a satisfactory inhibitory effect, therefore it was excluded from the studies. The mixtures were incubated for 30 minutes at RT. When the substrate with a final concentration of 50 μM was added to the mixture, the fluorescence intensities were measured at 60 s intervals over 30 minutes using an Infinite 200 PRO plate reader (Tecan, Männedorf, Switzerland). The temperature was set to 37 °C. The excitation and emission wavelengths were 360 nm and 460 nm, respectively. Inhibition assays were performed as triplicates.

For the quinacrine and suramin combination test, a 1:1 stock solution of the molecules was prepared and 0.5 μM of the protein was incubated with 0-75 μM of the combined molecules (quinacrine and suramin).

The IC_50_ value was calculated by plotting the initial velocity against various concentrations of the combined molecules using a dose-response curve in GraphPad Prism^5^ software. All measurements were performed in triplicate and data are presented as mean ± SD.

### 2.4 Determination of inhibition mode and inhibitory constant

The mode of inhibition was determined using different final concentrations of the inhibitors and substrate. Briefly, SARS-CoV-2 3CL^pro^ at 0.5 μM was incubated with the inhibitor at different concentrations for 30 minutes at RT. Subsequently, the reaction was initiated by the addition of the corresponding concentration series of the substrate. The data analysis was performed using a Lineweaver-Burk plot, therefore the reciprocal of velocity (1/V) vs. the reciprocal of the substrate concentration (1/[S]) was compared (Motulsky and Christopoulos, 2004; Roy et al., 2017). All measurements were performed in triplicate and data are presented as mean ± SD. The inhibitory constant (Ki) was obtained using equation (2):

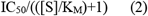

while IC_50_ and K_M_ were determined as described before and [S] is the concentration of substrate used in the experiments (Wu et al., 2014).

### 2.5 Determination of dissociation constant using fluorescence spectroscopy

The intrinsic Trp fluorescence of SARS-CoV-2 3CL^pro^ was measured under the influence of quinacrine and suramin, based on this experiment the dissociation constant between the protein and the ligands could be determined (Coronado et al., 2018). Briefly, protein sample was in 25 mM Tris-HCl pH 8.0, 150 mM NaCl with a final concentration of 10 μM/50 μl. The protein solution within the cuvette (1 cm path length) was titrated stepwise with a molecule stock solution 0-48 μM of quinacrine or suramin (to avoid protein dilution the molecule stock solution was treated with the same protein concentration - 10 μM protein + 500 μM molecule stock). A measurement was conducted following each titration. The quenching of the protease fluorescence, ΔF (F_max_-F), at 330 nm of each titration point was used for fitting a saturation binding curve using a nonlinear leastsquares fit procedure which has been discussed in detail elsewhere (Johnson and Frasier, 1985) based on equation (3) (Shaikh et al., 2007):

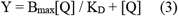

where, [Q] is the ligand concentration in solution, acting as a quencher, y is the specific binding derived by measuring fluorescence intensity, B_max_ is the maximum amount of the protease-ligand complex at saturation of the ligand and K_D_ is the equilibrium dissociation constant. The percentage of bound protease, i.e. y, derived from the fluorescence intensity maximum, is plotted against the ligand concentration. Additionally, the data were fitted with a modified Hill equation obtaining the following relation (4) (Wang et al., 2015; Ahumada et al., 2017):

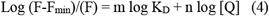

where, F_min_ is the minimal fluorescence intensity in the presence of the ligand, K_D_ is the equilibrium constant for the protein-ligand complex. The “binding constant” K is defined as the reciprocal of K_D_.

### 2.6 Circular dichroism (CD) spectroscopy

CD measurements were carried out with a Jasco J-1100 Spectropolarimeter (Jasco, Germany). Far-UV spectra were measured in 190 to 260 nm using a protein concentration of four μM in 20 mM K_2_HPO_4_/KH_2_PO_4_ pH 7.5. Cells of one mm path length were used for the measurements; 15 repeat scans were obtained for each sample and five scans were conducted to establish the respective baselines. The averaged baseline spectrum was subtracted from the averaged sample spectrum. The results are presented as molar ellipticity [θ], according to the equation (5):

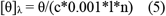

where θ is the ellipticity measured at the wavelength λ (deg), c is the protein concentration (mol/L), 0.001 the cell path length (cm) and n the number of amino acids. The results were analyzed and the secondary structure content was determined using the CDpro software package (Sreerama and Woody, 2000).

### 2.7 Systems information, molecular docking and ligand parameterization

The 3D model of SARS-CoV-2 3CL^pro^ was obtained from the PDB database (PDB entry: 6M2Q). The protonation state of the amino acid side chain was set to pH 7.4 using web-server H++ (Gordon et al., 2005), placed in a water box, neutralized and followed by a 200 ns molecular dynamics (MD) simulation and the representative structure was obtained by clustering analysis. Molecular docking was used to select the best pose for the ligand-protein interaction using Autodock Vina program (Trott and Olson, 2010). A grid box was defined for docking the molecule (quinacrine) near the active site and for docking in an allosteric site (suramin). These calculations were performed with the scoring function of the program that ranked several poses for each ligand. The best pose of each ligand was chosen to prepare the 3CL^pro^-quinacrine and -suramin complexes.

Ligands were parameterized using Gaussian 16 (Frisch et al., 2016) at the level of theory HF/6-31G* to optimize their geometry and calculate their electrostatic potentials. Restrained electrostatic potential (RESP) charges were determined using antechamber (Wang et al., 2006), while the general amber force field (GAFF) (Wang et al., 2004) was used for the missing parameters.

### 2.8 Simulation setup

All the molecular dynamics simulations were carried out using the Amber18 (Case et al., 2018) software package and the FF19SB (Tian et al., 2020) force field was used to describe the protein atoms interactions, while GAFF and RESP charges describe the quinacrine and suramin molecules. The systems were solvated in an octahedral box of TIP3P water molecules with at least 10 Å distance from any solvent atoms between the solute’s in each direction and the system was neutralized when necessary. Initially, energy minimization was performed in two steps to remove poor contacts from the initial structures. First, the protein/complex was constrained (force constant of 50.0 kcal/mol.Å^2^) and minimized using 5,000 steepest descent steps followed by 5,000 conjugate gradient steps and by a 10,000 steps unconstrained energy minimization round. The system was slowly heated from 0 to 298 K for 500 ps under constant atom number, volume and temperature (NVT) ensemble, while the protein was restrained with a force constant of 25 kcal/mol.Å^2^. After the heating process, the equilibrium stage was performed using constant atoms number, pressure and temperature (NpT) ensemble for 5 ns. Finally, the simulation run was performed for 200 ns (single protein) and 300 ns (complexes) without any restraints and under NVT ensemble and, the constant temperature and pressure (1 atm) were controlled by Langevin coupling. Long-range electrostatic interaction was calculated by the particle-mesh Ewald method (PME) (Darden et al., 1993) with 8 Å cutoff. The Shake constraints were applied to all bonds involving hydrogen atoms to allow a 2-fs dynamics time step.

### 2.9 Molecular dynamics analysis

The CPPTRAJ9 program of AmberToolsl9 (Case et al., 2005) was used to analyze the MD simulations. Root mean square deviation (RMSD) and the radius of gyration (Rg) of Cα were calculated to determine the system quality and stability and to determine the equilibration and convergence of the systems. Protein flexibility was calculated by the root mean square fluctuation (RMSF) for all Cα atoms, residue-by-residue over the equilibrated trajectories.

Clustering analysis was performed with the k-means method ranging from 2 to 6 and to access the quality of clustering the DBI values and silhouette analyses was used.

The interaction energy was calculated using the generalized Born (GB)-Neck2 (Nguyen et al., 2013) implicit solvent model (igb = 8). Molecular mechanics/generalized Born surface area (MM/GBSA) energy was computed between the protein and the ligand in a stable regime comprising the last 50 ns of the MD simulation, stripping all the solvent and ions. The web version of POCASA 1.1 was used to determine the volume of the active site pocket after MD simulations (Yu et al., 2010).

### 2.10 Statistical analysis

The significance of the inhibition assays and fluorescence spectroscopy experiments were evaluated by a one-way analysis of variance (ANOVA) followed by Bonferroni post hoc test. Differences were considered significant when a *P* value was less than 0.05 (Zhang et al., 1999) All statistical analyses were performed with GraphPad Prism software version 5 (San Diego, CA, USA).

## 3. Results and Discussion

### 3.1 Expression and Purification of SARS-CoV-2 3CL^pro^

SARS-CoV-2 3CL^pro^_GST fusion protein was expressed in *E. coli* Lemo21 (DE3) cells and purified using a GSH-Sepharose column **(Supplementary Fig. S1A)**. The relevant protein fractions were concentrated and prepared for PreScission protease cleavage to remove the GST-tag. The SDS gel **(Supplementary Fig. S1B and S1C)** indicate the cleavage efficiency and the purity of 3CL^pro^. The pure protein was concentrated and applied onto a Superdex 200 10/300 GL size exclusion chromatography (GE Healthcare) to remove aggregated protein species **(Supplementary Fig. S1D)**. CD spectroscopy of 3CL^pro^ indicated that the protein is correctly folded after purification process and GST-cleavage. Deconvolution of the CD data using the software CDpro (Sreerama and Woody, 2000) showed that the protein secondary structure contains around 28% α-helices and 23% β-strands. Which is in good agreements with data from the crystal structure (PDB entry: 6M2Q) which contain 25% α-helices and 28% β-strands **(Supplementary Fig. S1E)**.

### 3.2 Activity assay of SARS-CoV-2 3CL^pro^

The SARS-CoV-2 3CL^pro^ activity was investigated using an assay procedure described earlier (Zhang et al., 2020a; Zhang et al., 2020b; Ma et al., 2020) and DABCYL-KTSAVLQ/SGFRKME-EDANS (Bachem, Switzerland) was used as substrate. Activity assays were performed to obtain the kinetic key values V_max_, K_M_ and K_cat_. A standard curve was generated converting the relative fluorescence unit (RFU) to the amount of cleaved substrate (μM) **(Supplementary Fig. S2A)**. In the next step, enzymatic activity of the protease was characterized by measuring V_max_ and K_M_ values; 0.5 μM of the protease was mixed with various concentrations of the substrate (0-200 μM). The initial velocity was measured and plotted against the substrate concentration. Curve fitting with Michaelis-Menten equation resulted in the best fitting values of K_M_ and V_max_ as 25.47 ± 3.43 μM and 47.52 ± 2.91 μM/s, respectively **(Supplementary Fig. S2B)**. The calculated K_cat_/K_M_ was 3,731.21 s^−1^ M^−1^ which is like the previously reported values 3,4261.1 s^−1^ M^−1^ and 5,624 s^−1^ M^−1^ (Zhang et al., 2020a; Ma et al., 2020).

### 3.3 Inhibition assay of chloroquine, quinacrine and suramin against SARS-CoV-2 3CL^pro^

A primary inhibition test of the three antiparasitic compounds chloroquine, quinacrine and suramin (10 μM) was performed against SARS-CoV-2 3CL^pro^ to screen the best inhibitor against the virus protease (**Fig. 1**).

**Fig. 1.**
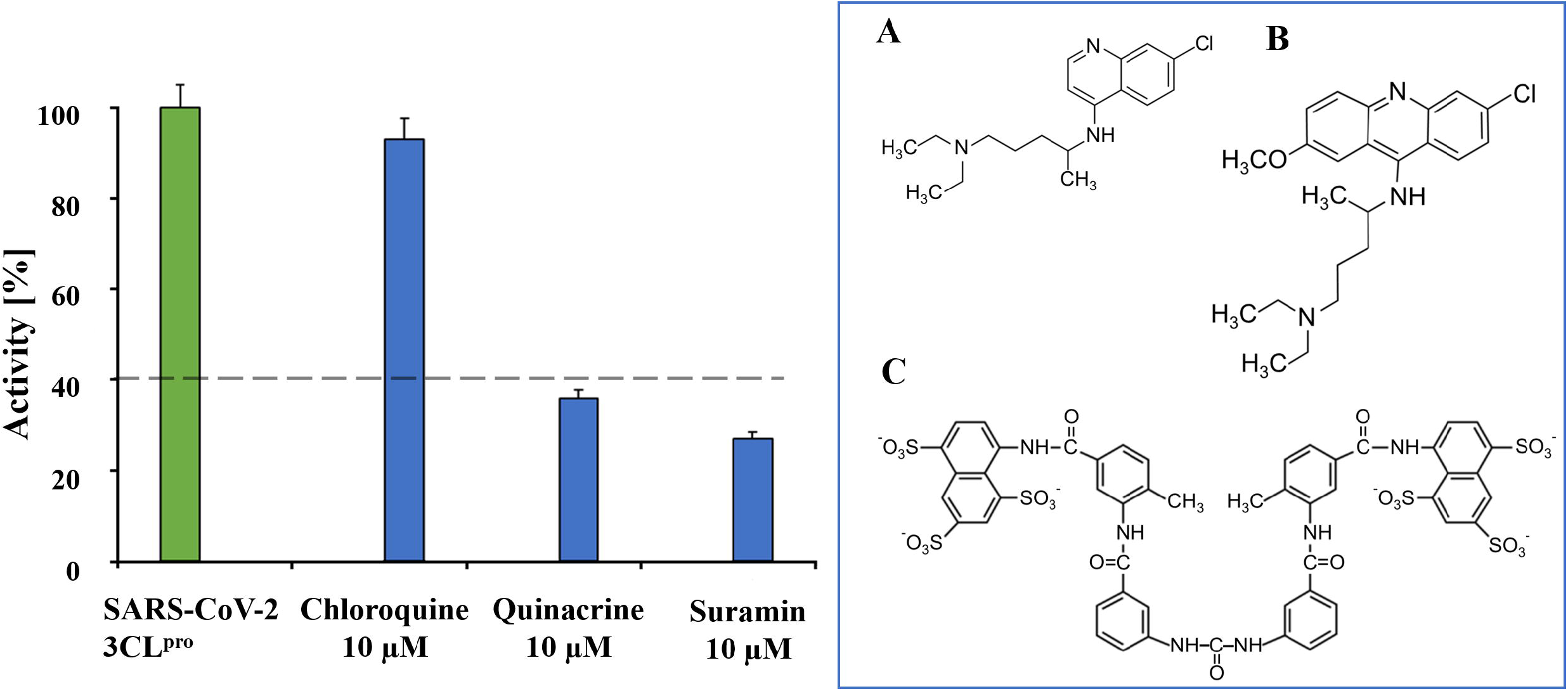
Preliminary inhibition tests of chloroquine, quinacrine and suramin against SARS-CoV-2 3CL^pro^. Quinacrine and suramin inhibit the virus protease activity by more than 60%. Contrary, chloroquine shows not relevant inhibition, just about 10%. The blue box shows the molecular structures of **(A)** Chloroquine, **(B)** Quinacrine and **(C)** Suramin.

The primary inhibition tests showed a strong effect of quinacrine and suramin against SARS-CoV-2 3CL^pro^ activity. In contrast, chloroquine has a weak effect on 3CL^pro^ proteolytic activity. Suramin and quinacrine possess an anti-viral effect against SARS-CoV-2 in cell culture (da Silva et al., 2020; Duan et al., 2020; Roldan et al., 2020; Touret and de Lamballerie, 2020), however, the target protein of both molecules in the viral replication process was not identified. Based on the primary inhibition results, quinacrine and suramin were further investigated regarding their inhibitory potential, and our results demonstrate that suramin and quinacrine have an inhibitory effect against the 3CL protease of SARS-CoV-2 **(Figures 2 and 3, respectively)**.

**Fig. 2.**
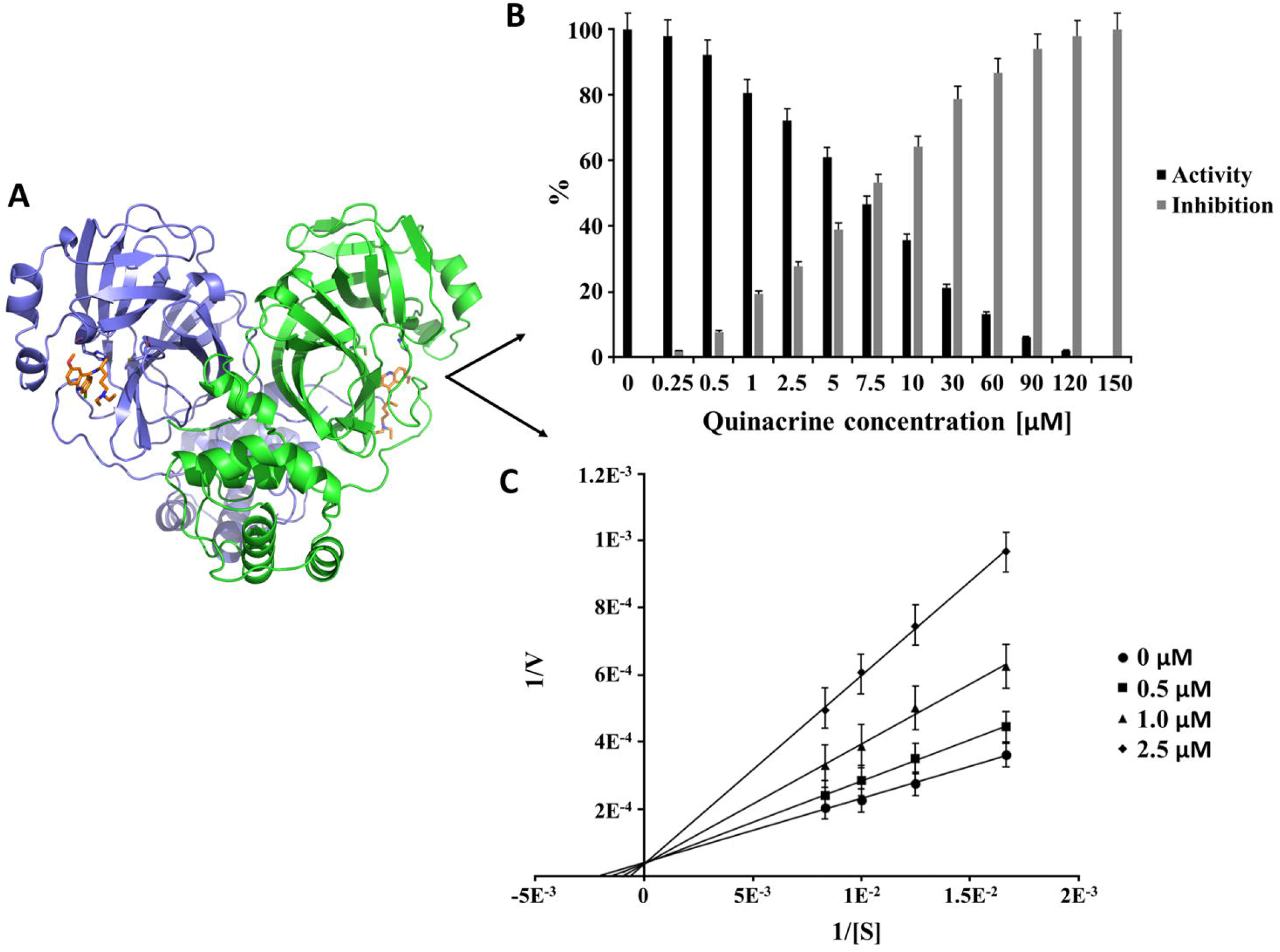
Inhibition effect and inhibition mode of quinacrine over SARS-CoV-2 3CL^pro^. **(A)** 3D representation of the 3CL^pro^-quinacrine complex (3CL^pro^ in Ribbon and quinacrine in Sticks). 3CL^pro^ monomers are colored in green and blue, and quinacrine in orange. Normalized activity and inhibition of the virus proteases and, Lineweaver-Burk plots to determine the inhibition mode are presented. [S] is the substrate concentration; v is the initial reaction rate. **(B)** Normalized activity and inhibition of SARS-CoV-2 3CL^pro^ under quinacrine influence. **(C)** Lineweaver-Burk plot for quinacrine inhibition of SARS-CoV-2 3CL^pro^.

Quinacrine inhibit the protease activity 100% at a concentration of 150 μM **(Fig. 2B)** and suramin at 100 μM **(Fig. 3B)**. Quinacrine has a calculated IC_50_ value of 7.8 ± 0.9 μM **(Table 1 and Supplementary Fig. S3A)**. Further experiments identified quinacrine as a competitive inhibitor of the 3CL^pro^ **(Fig. 2C)**, it means that the molecule interacts directly with amino acid residues located in the active site and/or with the substrate-binding site. In contrast, suramin acts as a non-competitive inhibitor **(Fig. 3)**, showing a type of allosteric inhibition.

**Fig. 3.**
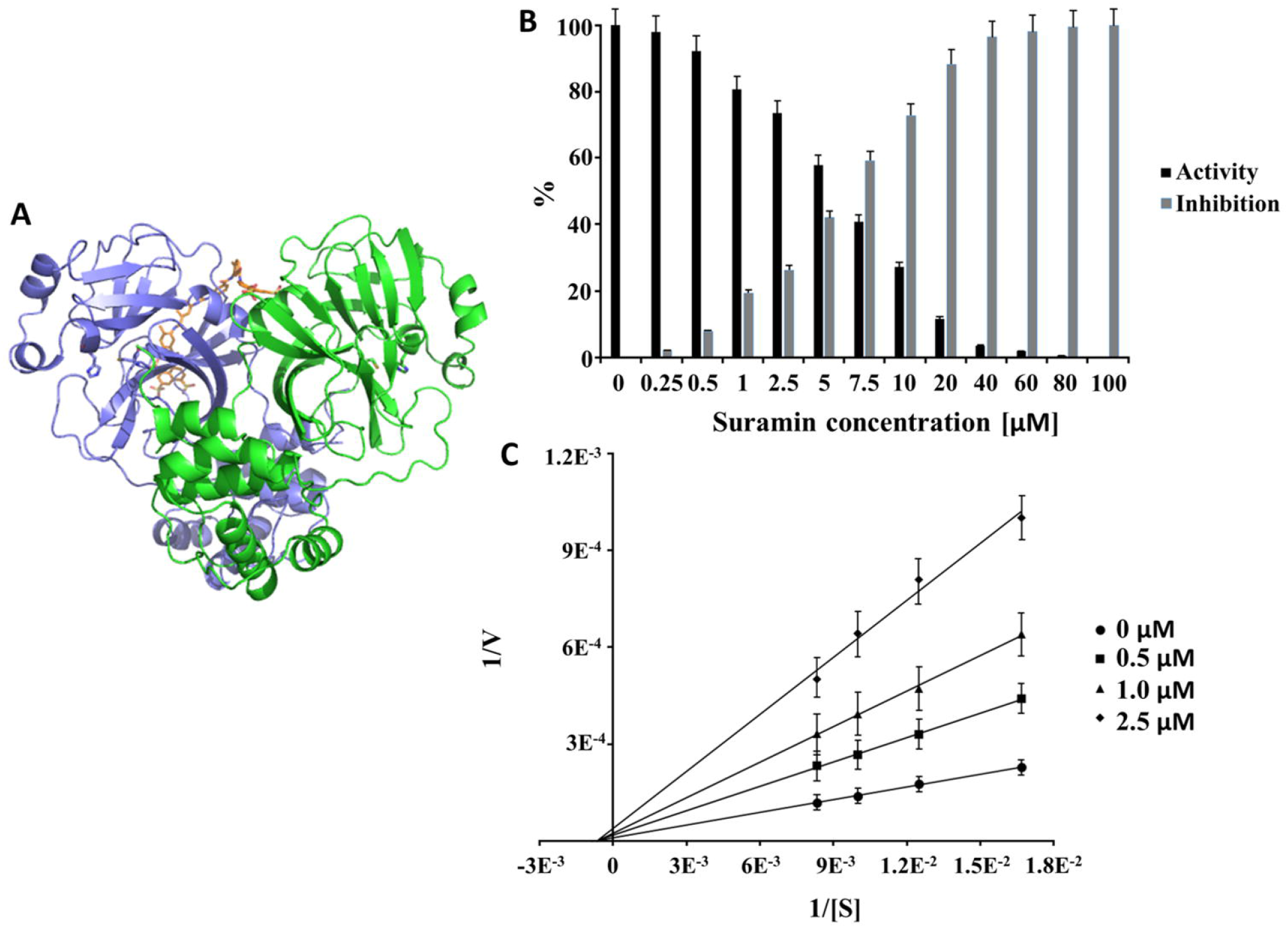
Inhibition effect and inhibition mode of suramin over SARS-CoV-2 3CL^pro^. **(A)** 3D representation of the 3CL^pro^-suramin complex (3CL^pro^ in Ribbon and suramin in Sticks). 3CL^pro^ monomers are colored in green and blue and suramin in orange. Normalized activity and inhibition of the virus proteases and, Lineweaver-Burk plots to determine the inhibition mode are presented. [S] is the substrate concentration; v is the initial reaction rate. **(B)** Normalized activity and inhibition of SARS-CoV-2 3CL^pro^ under suramin influence. **(C)** Lineweaver-Burk plot for suramin inhibition of SARS-CoV-2 3CL^pro^.

**Table 1.** Summary of the SARS-CoV-2 3CL^pro^ inhibition experiments by quinacrine and suramin.

| Molecule | IC<sub>50</sub> [$\mu\text{M}$] | K<sub>i</sub> [$\mu\text{M}$] | Inhibition type |
| --- | --- | --- | --- |
| Quinacrine | $7.8 \pm 0.9$ | $1.6 \pm 0.4$ | competitive |
| Suramin | $6.3 \pm 0.8$ | $1.3 \pm 0.4$ | non-competitive |

The potency of suramin in inhibiting 3CL^pro^ was calculated and demonstrated an IC_50_ value of 6.3 ± 0.8 μM **(Table 1 and Supplementary Fig. S3B)**.

The IC_50_ values of both molecules are in a concentration range with other repurposed drug molecules, which also demonstrate inhibitory effect against SARS-CoV-2 3CL^pro^, for example; boceprevir has an IC_50_ value of 4.14 μM (Ma et al., 2020), menadione presents IC_50_ value of 7.9 μM (He et al., 2020) and disulfiram has an IC_50_ of 9.35 μM (Jin et al., 2020).

### 3.4 Investigation of SARS-CoV-2 3CL^pro^ with suramin and quinacrine using fluorescence spectroscopy

To confirm the viability of the assay, fluorescence spectra from the intrinsic tryptophan (Trp) of the SARS-CoV-2 3CL^pro^ with the inhibitor molecules were investigated. The fluorescence changes of Trp can reflect environmental variation of the protein and the decreased Trp emission intensity confirmed the complex formation between the protease and the inhibitory molecules. Based on the quenching of the protease fluorescence of each titration point, the dissociation constant (KD) could be determined for each molecule (suramin and quinacrine) **(Supplementary Fig. S4)**.

**Table 2** describe the K_D_ value of the suramin interaction with SARS-CoV-2 3CL^pro^ (4.5 ± 1.2 μM) with slight increase for quinacrine (6.0 ± 1.8 μM). Interestingly, the protein interaction with suramin or quinacrine induce a different type of excitation shift of the protein tryptophans. Suramin interaction induces a conformational change in the protein structure, which was demonstrated by a red edge excitation shift (REES) of about 30 nm (330 to 360 nm) **(Supplementary Fig. S4A)**. REES is defined through increasing interactions between the fluorophore (Trp) and the surrounding solvent in the ground and excited state (Catici et al., 2016). On the contrary, the quinacrine interaction induced a blue edge excitation shift (BEES) of around 10 nm (330 to 320 nm) **(Supplementary Fig. S4C)**. Through BEES, the environment hydrophobicity of the Trp residue increases significantly (Möller and Denicola, 2002).

**Table 2.** Summary of the SARS-CoV-2 3CL^pro^ in complex with quinacrine and suramin.

| Molecule | K<sub>D</sub> [μM] | Excitation shift |
| --- | --- | --- |
| Quinacrine | 6.0 ± 1.8 | blue edge excitation shift |
| Suramin | 4.5 ± 1.2 | red edge excitation shift |

### 3.5 Docking and molecular dynamic simulations of suramin and quinacrine with SARS-CoV-2 3CL^pro^ structure

The atomic coordinates from SARS-CoV-2 3CL^pro^ (PDB entry: 6M2Q) was used as initial model. Cluster analysis of the MD simulations demonstrated that cluster #0 showed the best results with representativeness of over 86.1% of the whole simulation time and the selected structure appeared during the simulation around 169 ns **(Supplementary Fig. S5)**. SARS-CoV-2 3CL^pro^ is a homodimer with the catalytical dyad His41 and Cys145 (Zhang et al., 2020a). Dimerization of the protease is necessary for catalytic activity; the interaction of each protomer N-terminal Ser1 with Glu166 of the other protomer stabilizes the shape of the S1 pocket, which is important for the binding of the substrate (Anand et al., 2002). Chen et al. 2006 suggested that only one protomer of the SARS-CoV 3CL^pro^ dimer is active (Chen et al., 2006). Interestingly, after the initial MD simulations of SARS-CoV-2 3CL^pro^ dimer **(Supplementary Fig. S6)**, one protomer active site collapsed and, the volume of the pocket decreased by around twofold, from 261 to 142 Å^3^ compared to the active site of the other protomer. The main difference observed between the two protomers succeeding the MD simulation was the interaction between the C-terminal Gln306 and Ile152 of the same protomer. A hydrogen bond was observed between NE2 of Gln306 and the backbone oxygen of Ile152 **(Supplementary Fig. S7)**. We assume that this interaction in addition to the already described (Glu166 with Ser1) stabilizes the active site. To conceive the molecular binding modes of the inhibitory molecules, theoretical investigations through docking and molecular dynamics simulations were carried out. The interactions between SARS-CoV-2 3CL^pro^ in complex with quinacrine or suramin were analyzed to predict their binding mode. Quinacrine was docked in both protease active sites (as competitive inhibitor) and suramin (as non-competitive inhibitor) in a region with strong positive charges on the protein surface, as suramin is a highly negatively charged molecule (Burch and Ashburn, 1951). Subsequent MD simulations demonstrated a stable system for a period of 300 ns **(Supplementary Fig. S8 and S9)**. The binding assessment of the ligands to the protease was observed using a residue-wise decomposition of the binding energy received from the MD simulations. **Table 3 and Fig. 4** illustrate the 3CL^pro^ amino acid residues contribution to the binding energy in the active site that interact with quinacrine. The interactions between quinacrine and the protease are stabilized by two hydrogen bonds (H-bond), which are mediated by donor and acceptor atoms of Met165 and Gln189 **(Table 4)**. His41, Met49, Val186, Arg188 and Gln194 interact with quinacrine by hydrophobic interactions **(Table 3 and Fig. 4)**.

**Fig. 4.**
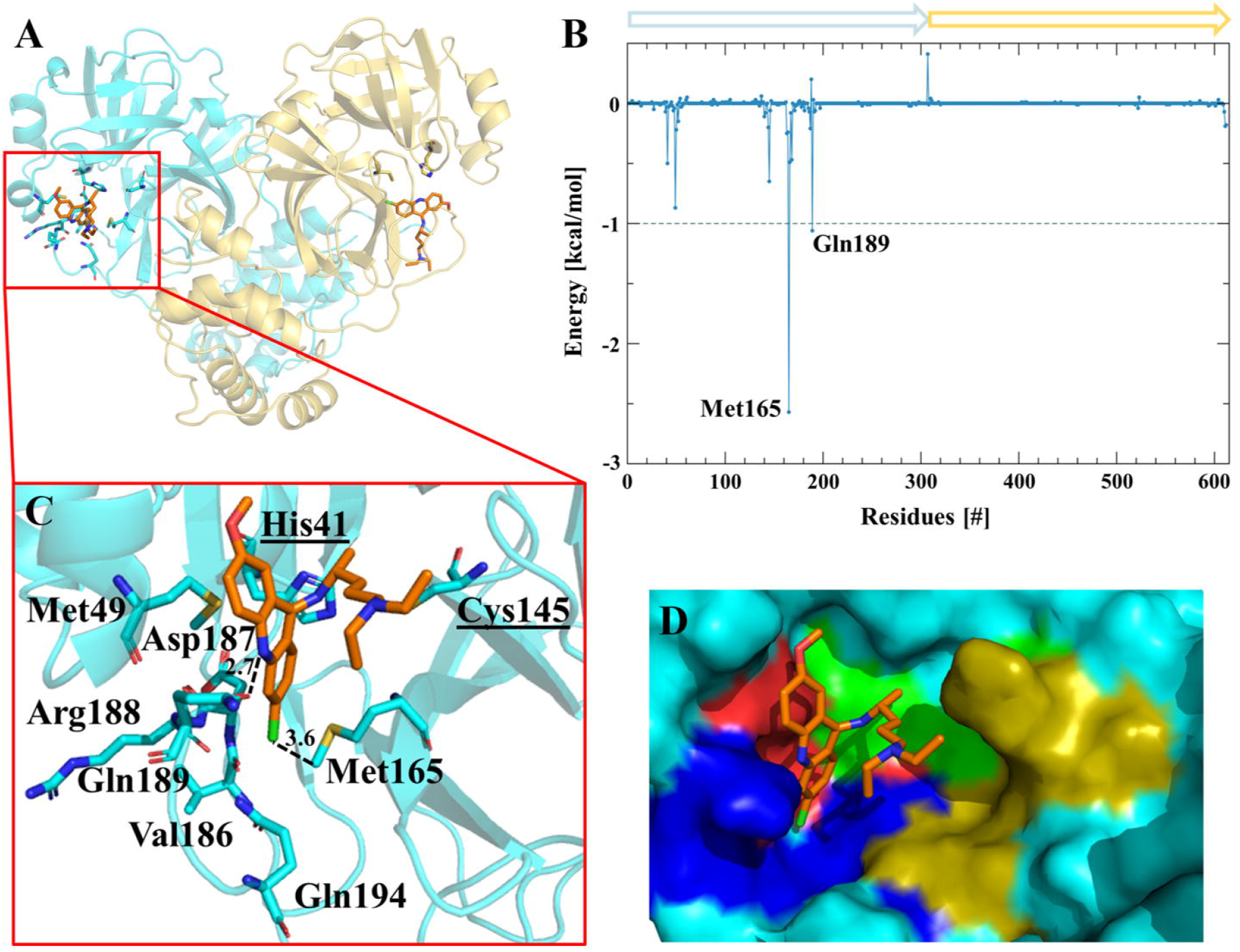
Amino acids participating in the SARS-CoV-2 3CL^pro^-quinacrine interaction. **(A)** Ribbon and sticks representation of the SARS-CoV-2 3CL^pro^-quinacrine complex after MD simulation. **(B)** Decomposition of the binding energy of SARS-CoV-2 3CL^pro^-quinacrine complex. Arrows label protomer A (turquoise) and protomer B (gold). **(C)** SARS-CoV-2 3CL^pro^ amino acids involved in the interaction with quinacrine based on MD simulations. **(D)** Quinacrine interaction in the protease substrate-binding and active site. The substrate-binding subsites are highlighted: green S1’: gold S1; red S2 and blue S3.

**Table 3.** SARS-CoV-2 3CL^pro^ Residues involved in forming H-bonds and hydrophobic contacts with quinacrine and suramin.

| Ligand | Interacting Residues |  |
| --- | --- | --- |
| Quinacrine | Met165, Gln189 | His41, Met49, Val186, Asp187, Arg188, Gln194 |
| Suramin | Lys12, Lys97, Lys100, Tyr101, Phe103 | Lys97, Lys100, Lys102, Phe103, Val104, Arg105 |

**Table 4.**
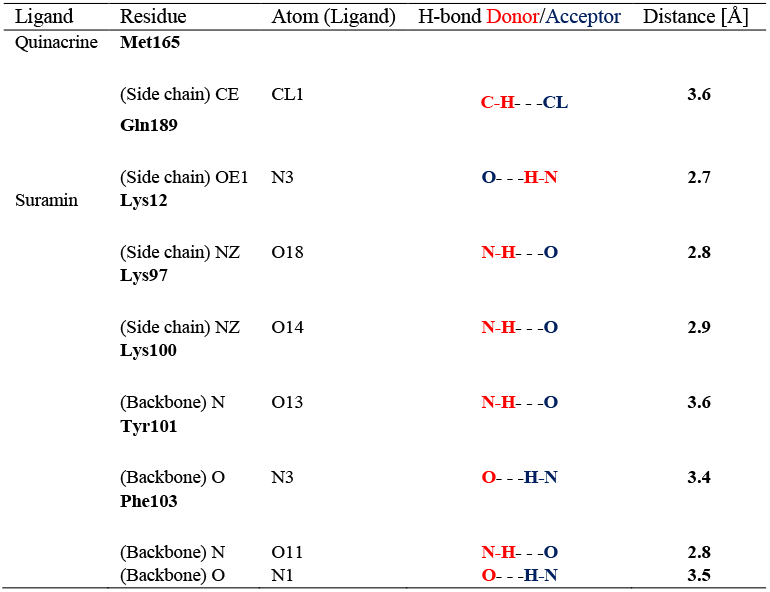
Atoms involved in the H-Bond interaction between SARS-CoV-2 3CL^pro^ and inhibitors.

**Fig. 5** shows the amino acid residues involved in the interaction between 3CL^pro^ and suramin, **Tables 3 and 4** summarize the H-bonds and hydrophobic interactions. The interactions that stabilize the protease-suramin complex are mediated by six H-bonds that are formed by Lys12, Lys97, Lys100, Tyr101 and Phe103. Additionally, the ligand is stabilized by hydrophobic interactions with the residues: Lys97, Lys100, Lys102, Phe103, Val104 and Arg105 (**Table 3**).

**Fig. 5.**
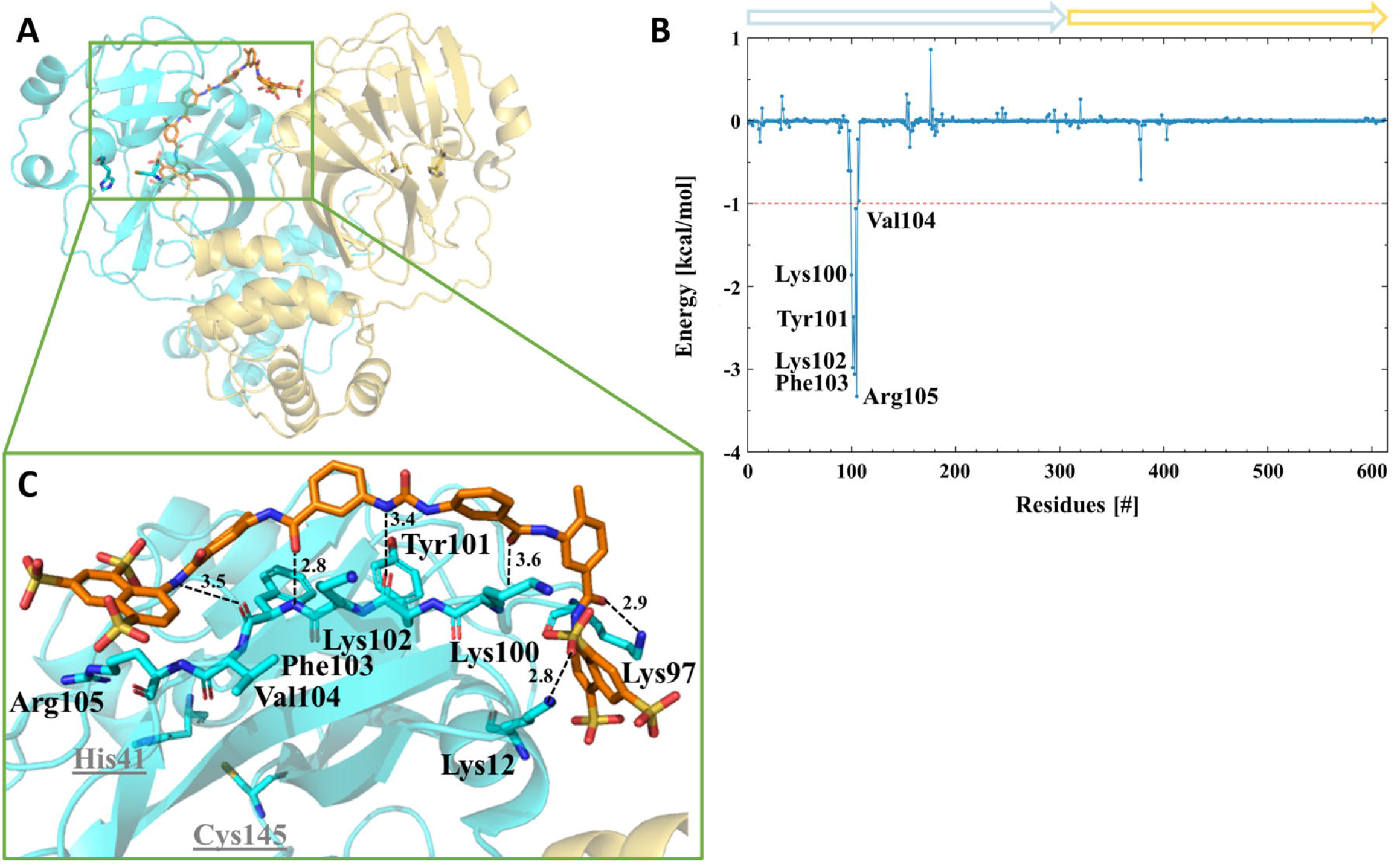
Amino acids participating in the SARS-CoV-2 3CL^pro^-suramin interaction. **(A)** Ribbon and sticks representation of the SARS-CoV-2 3CL^pro^-Suramin complex after MD simulation. **(B)** Decomposition of the binding energy of SARS-CoV-2 3CL^pro^-suramin complex. Arrows label protomer A (turquoise) and protomer B (gold). **(C)** SARS-CoV-2 3CL^pro^ amino acids involved in the interaction with Suramin based on MD simulations.

As a competitive inhibitor of SARS-CoV-2 3CL^pro^, quinacrine interacts through hydrophobic interaction with His41 of the protease catalytic dyad, blocking the entrance to the active site.

On the contrary, suramin is not directly interacting with the active site residues and our results suggested that suramin interact with the protein allosterically changing the catalytic site conformation, thus preventing the substrate entry.

An important task is the suitability of the inhibitor to the binding areas so that covalent inhibition occurs (Pettinger et al., 2017). Therefore, the nucleophilicity of the amino acid residues in the target and electrophilic groups in the drug need to be considered (Way, 2000). The relative nucleophilicity of the amino acid residues in their neutral states are given in the order of Cys (1) > His (10^−2^) > Met (10^−3^) > Lys, Ser (10^−5^) > Thr and Tyr (10^−6^) (Way, 2000; Pettinger et al., 2017). This is one of the reasons why many of the covalent modifiers in drugs are designed to target the thiol group of cysteine (Gan et al., 2009; Leproult et al., 2011). Quinacrine interact directly with nucleophilic residues in the active site of SARS-CoV-2 3CL^pro^ (e.g. His41 and Met165) which make them highly attractive for covalent interactions. Contrary, suramin interact with Lys and Tyr residues that can also form covalent bonds (Tang et al., 2020), but in a less probability compared with Cys, His and Met.

To study in more detail the plasticity of the virus protease in complex with quinacrine and suramin the consequences on the ligand binding areas, the substrate-binding sites (S1, S1’, S2, S3) and the oxyanion hole after MD simulation of the complexes were analysed. Amino acid residues forming the substrate-binding site of SARS-CoV-2 3CL^pro^ and their subsites were described previously **(Supplementary Table S1)** (Chen et al., 2006; Tang et al., 2020). **Fig. 6** describe the changes occurred on SARS-CoV-2 3CL^pro^ substrate-binding site caused by the interaction of the inhibitors. MD simulations revealed possible mode of interaction of quinacrine with SARS-CoV-2 3CL^pro^. Thereby, His41, one amino acid residue of the catalytic dyad, interact with quinacrine, this interaction is responsible to block partially the S1’ and S3 sites occupying completely the S2 subsite **(Fig. 6)**. Four residues of the protease S2 subsite interact directly with quinacrine via hydrophobic interactions (His41, Met49, Asp187 and Arg188) and two residues of the S3 subsite form H-bonds with the inhibitor molecule (Met165 and Gln189). In the case of suramin, no single amino acid residue of the S1, S1’, S2 and S3 subsites are in interaction with the ligand, which agrees with the results of the activity assay that describe suramin as non-competitive inhibitor of SARS-CoV-2 3CL^pro^.

**Fig. 6.**
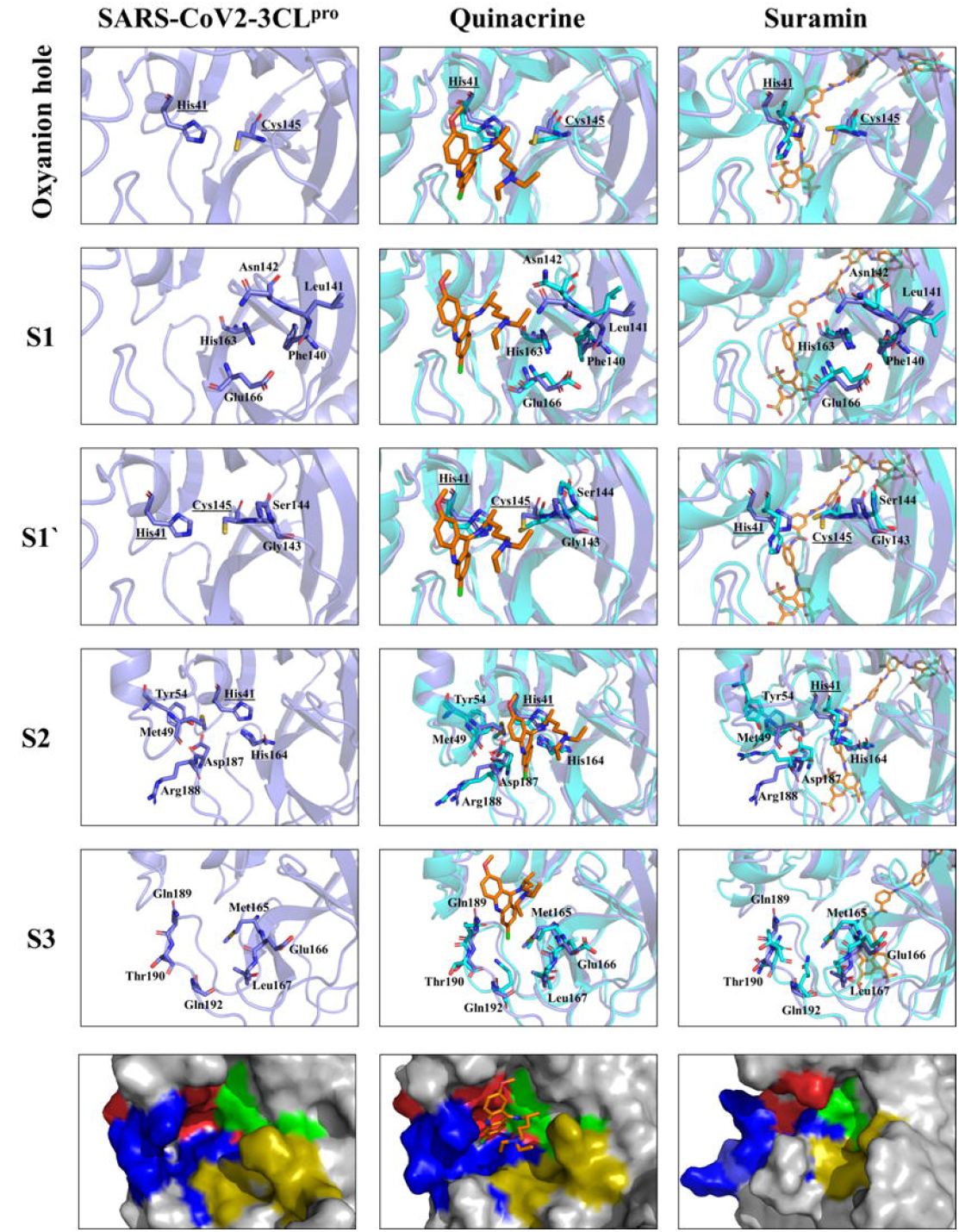
Substrate-binding pocket and oxyanion hole of SARS-CoV-2 3CL^pro^ (S1, S1’, S2, S3) free accessibility and under the influence of quinacrine and suramin. Structural overlay between the single protein and in complex, the active site residues are highlighted. The amino acid residues are shown in sticks. The surface view (zoom) of the substrate-binding area demonstrates the occupied area by the molecules and the conformational changes induced by its binding. Substrate-binding subsites highlighted as green S1’; gold S1; red S2 and blue S3.

Suramin binds allosterically and the ligand interaction induces conformational changes in the catalytic site (most observable for subsites S1’ and S2), thus preventing the entry and the turnover of the substrate **(Fig. 7)**. Additionally, the volume of the active site pocket decreased considerably after suramin binding, even whether the molecule was docking just in protomer 1, changing the volume of the pocket from 261 to 90 Å^3^ (Protomer 1) and to 120 (Protomer 2) **(Fig. 7C to E)**.

**Fig. 7.**
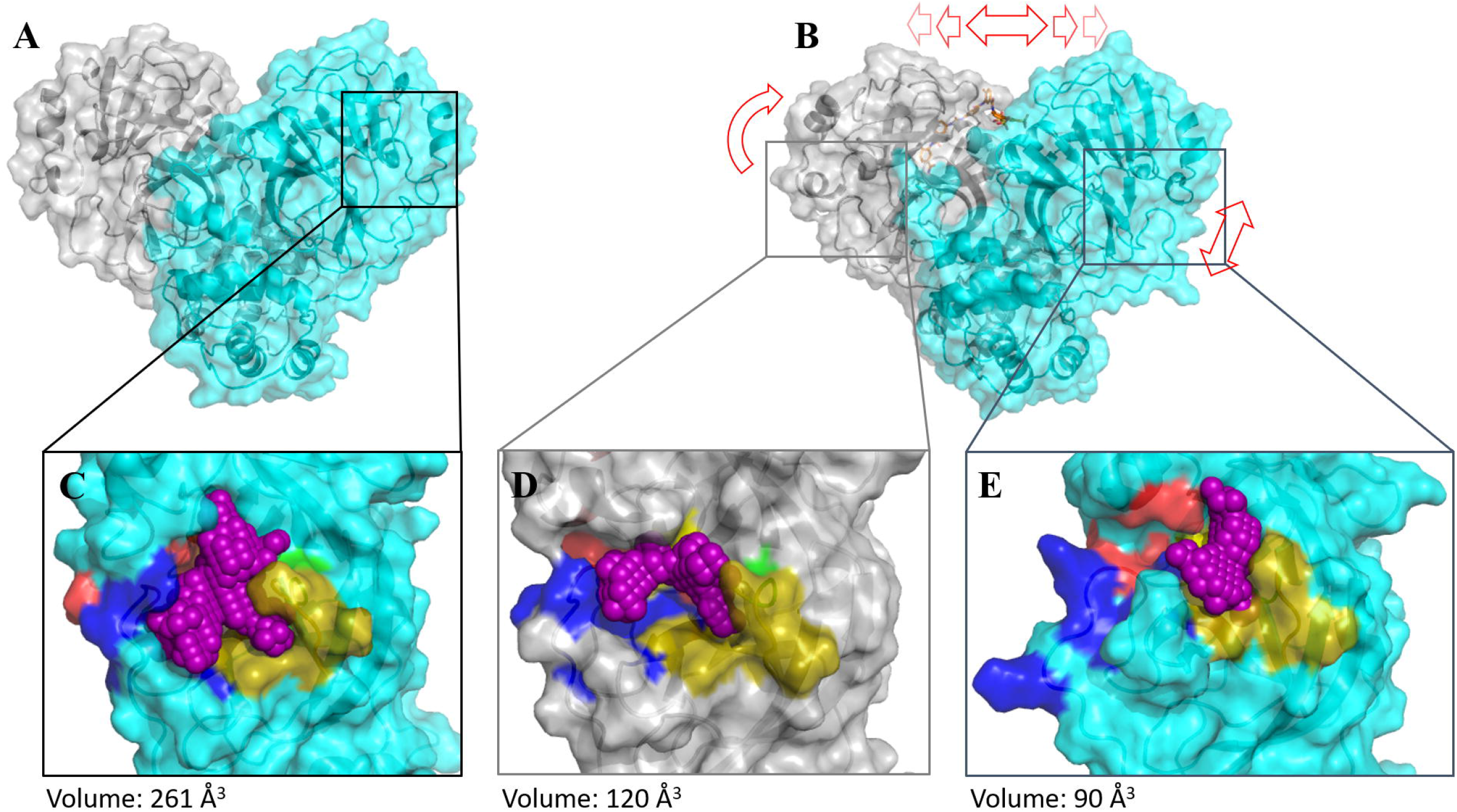
Comparison of SARS-CoV-2 3CL^pro^ with and without suramin after MD simulation. Protomer 1 is colored in turquois, protomer 2 in grey and active site pocket volume in violet. Red arrows highlight a conformational change in the SARS-CoV-2 3CL^pro^/suramin complex. **(A)** SARS-CoV-2 3CL^pro^ dimer. **(B)** SARS-CoV-2 3CL^pro^/suramin complex. **(C)** Active site pocket volume of SARS-CoV-2 3CL^pro^, substrate-binding subsites are highlighted as green S1’; gold S1; red S2 and blue S3. **(D)** Active site pocket volume of protomer 2. **(E)** Active site pocket volume of protomer 1.

### 3.6 Suramin and quinacrine act cooperatively to inhibit SARS-CoV-2 3CL^pro^

Suramin and quinacrine, as described previously, bind in different regions of SARS-CoV-2 3CL^pro^.To conjecture about the effect of both molecules simultaneously, we performed a combined inhibitory assay, whither both molecules were mixed (1:1) and tested against SARS-CoV-2 3CL^pro^ **(Fig. 8)**.

**Fig. 8.**
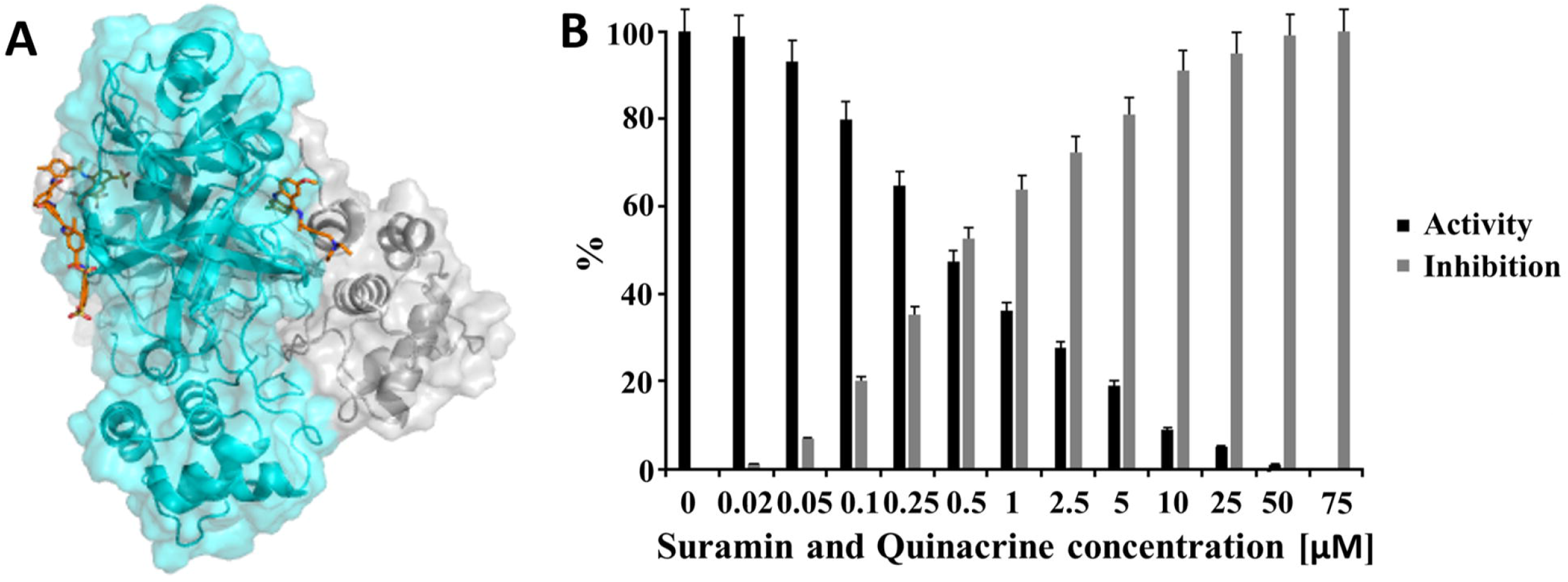
SARS-CoV-2 3CL^pro^ inhibition by a combination of quinacrine and suramin. **(A)** Ribbon representation of the SARS-CoV-2 3CL^pro^ model (grey/cyan) (PDB entry: 6M2Q) in complex with suramin and quinacrine (sticks in orange). **(B)** Normalized activity and inhibition of SARS-CoV-2 3CL^pro^ under quinacrine and suramin influence.

The inhibition assay was performed with a solution containing both ligands (suramin and quinacrine) with the molar ratio of 1:1. The calculated IC_50_ value (0.46 ± 0.1 μM) demonstrated an increased inhibition capacity of the drugs **(Table 5 and supplementary Fig. S11)**. When compared with the single molecules, the combination enhances the inhibitory capacity of the studied drugs for approx. 10 times. Which could be observed by the determined IC_50_ values. We assume that the conformational changes after suramin interaction with SARS-CoV-2 3CL^pro^ induce a structural change in both active sites in the dimer, which could make them more accessible for the interaction with quinacrine **(Fig. 7)**.

**Table 5.** Examples of drugs or preclinical tested compounds

| Compound | IC<sub>50</sub> [μM] | Reference |
| --- | --- | --- |
| Quinacrine | 7.8 ± 0.9 | reported here |
| Suramin | 6.3 ± 0.8 | reported here |
| Quinacrine + Suramin | 0.46 ± 0.1 | reported here |
| Boceprevir | 4.14 ± 0.61 | Ma et al., 2020 |
| Narlaprevir | 5.73 ± 0.67 | Ma et al., 2020 |
| Simeprevir | 13.74 ± 3.75 | Ma et al., 2020 |
| Auranofin | 0.51* | He et al., 2020 |
| Menadion | 7.96* | He et al., 2020 |
| Ebselen | 0.67 ± 0.06 | Jin et al., 2020 |
| Disulfiram | 9.35 ± 0.18 | Jin et al., 2020 |
| Tideglusib | 1.55 ± 0.30 | Jin et al., 2020 |
| Carmofur | 1.82 ± 0.06 | Jin et al., 2020 |

\* He et al. 2020 without standard deviation.

## 4. Conclusions

In summary, the cooperative inhibition of quinacrine and suramin could lead to a possible alternative to combat COVID-19 infections. Repurposing of clinically developed drugs is a common procedure to combat fast evolving pathogens. Similar studies addressed the use of established drugs against SARS-CoV-2 (e.g. Boceprevir, Menadione and Disulfiram) (Ma et al., 2020; He et al., 2020; Jin et al., 2020). **(Table 5)**. The 1:1 combination of quinacrine and suramin possess an effective anti-3CL^pro^ activity, demonstrating the potential of repurposing drugs to stop the replication process of SARS-CoV-2.

## Acknowledgements

We would like to thank members of the Multiuser Center for Biomolecular Innovation (IBILCE, São Jose do Rio Preto, Brazil) and the Institute of Biological Information Processing (Forschungszentrum Jülich, Germany), without whose help this project would have not been feasible. This research was supported by grants from CNPq [Grant numbers 435913/2016-6, 401270/2014-9, 307338/2014-2], FAPESP [Grant numbers 2016/12904-0, 2018/12659-0 2018/07572-3, 2019/05614-3], FUNDECT [23/200.307/2014], MOI III, CAPES and PROPe UNESP. This study was supported in part by the UFMS/MEC-Brasil and UFT/MEC-Brasil.

## Authors’ contributions

M.A.C. and R.J.E conceived, designed, performed and analyzed the experiments of SARS-CoV-2 3CL^pro^ in complex with Quinacrine and Suramin. D.S.O. and M.S.A. performed the docking and MD simulations. D.S.O., M.A.C, and R.J.E performed the docking and MD simulation analysis. R.K.A. and D.W. guided the research and the elaboration of the manuscript. R.J.E. and M.A.C. coordinated the research and wrote the manuscript. All authors read and took part in revising the final manuscript version.

