## Supplementary Material for "The repurposed drugs suramin and quinacrine inhibit cooperatively *in vitro* SARS-CoV-2 3CL^pro^"

Raphael J. Eberle^1,2#^*, Danilo S. Olivier^3^, Marcos S. Amaral^4^, Dieter Willbold^2,5,6^, Raghuvir K. Arni^1^, Monika A. Coronado^1,2#^*

^1^Multiuser Center for Biomolecular Innovation, IBILCE, Universidade Estadual Paulista (UNESP), São Jose do Rio Preto-SP, Brazil.

^2^Institute of Biological Information Processing (IBI-7: Structural Biochemistry), Forschungszentrum Jülich, Jülich, Germany.

^3^Federal University of Tocantins, Araguaína-TO, Brazil.

^4^Institute of Physics, Federal University of Mato Grosso do Sul, Campo Grande-MS, Brazil.

^5^Institut für Physikalische Biologie, Heinrich-Heine-Universität Düsseldorf, Universitätsstraße, Düsseldorf, Germany.

^6^JuStruct: Jülich Centre for Structural Biology, Forchungszentrum Jülich, Jülich, Germany.

^#^both authors contributed equally

**Table of contents**

Supplementary Figure S1. Purification of SARS-Cov-2 3CL^pro^ and quality check of the produced protein.

Supplementary Figure S2. K_M_ determination of SARS-CoV-2 3CL^pro^.

Supplementary Figure S3. Dose response curves for IC_50_ determination.

Supplementary Figure S4. Fluorescence spectroscopy of Trp at 295 nm of SARS-CoV-2 3CL^pro^ in the presence of the suramin and quinacrine.

Supplementary Figure S5. Clustering analysis of the SARS-CoV-2 3CL^pro^ structure after MD simulation.

Supplementary Figure S6. Time dependent modifications of the SARS-CoV-2 3CL^pro^ dimer.

Supplementary Figure S7. Time dependent modifications of SARS-CoV-2 3CL^pro^ dimer in the presence of quinacrine in each active site.

Supplementary Figure S8. Time dependent modifications of SARS-CoV-2 3CLpro dimer in the presence of suramin. Supplementary Figure S9. Atom numbers of quinacrine and suramin.

Supplementary Figure S10. Dose response curve for IC_50_ determination.

Supplementary Figure S11. SARS-CoV2 3CLpro dimeric structure after MD simulation.

Table S1. Contributing amino acids in the substrate-binding sites of SARS-CoV-2 3CL^pro^.

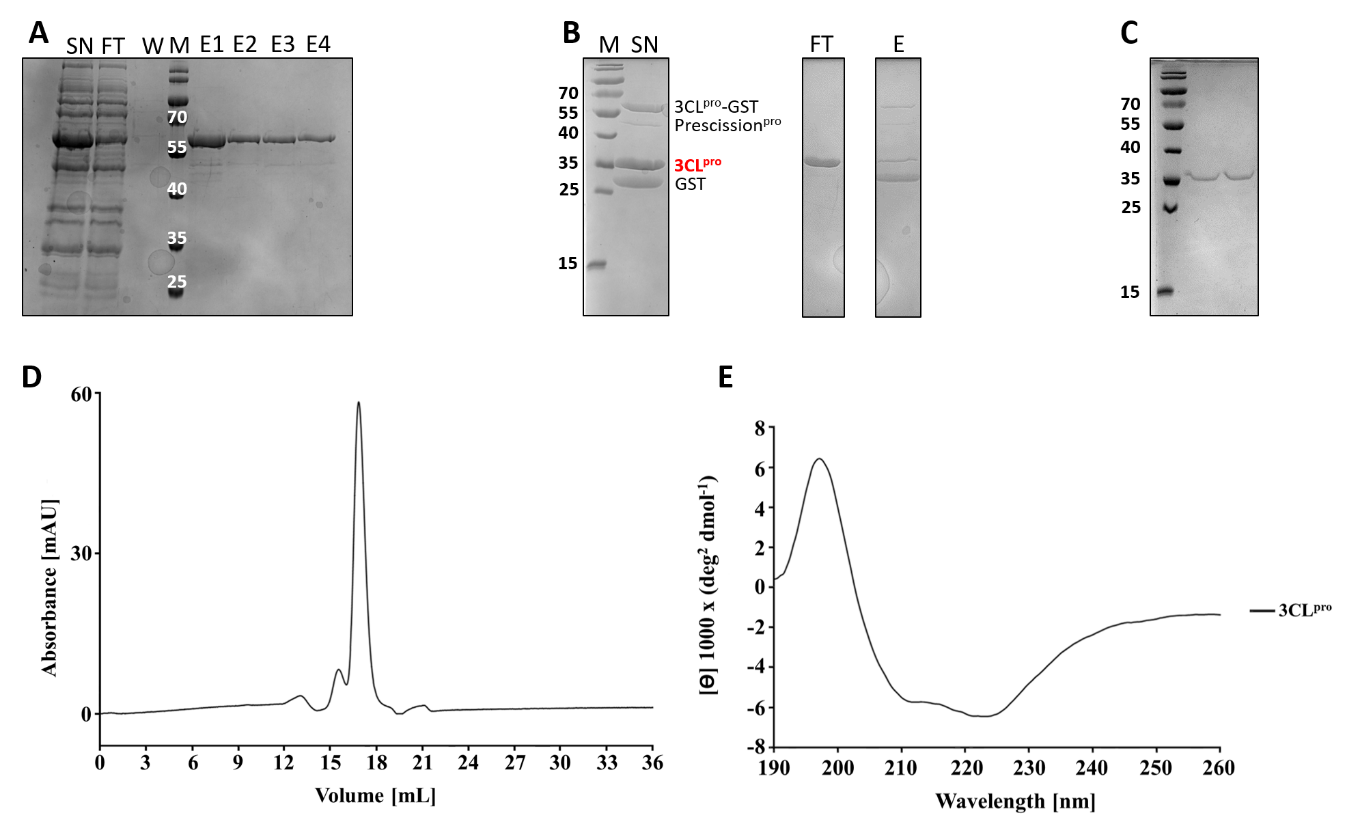

**Figure S1 Purification of SARS-Cov-2 3CL^pro^ and quality check of the produced protein.** **(A)** SDS 15% Gel of 3CL^pro^_GST purification with GSH sepharose. SN: supernatant, FT: flow through, W: washing step, M: protein marker, E1-E4: Elution. **(B)** SDS 15% Gel of the 3CL^pro^_GST fusion protein cleavage by PreScission^pro^. M: protein marker. SN: cleaving approach, containing 3CL^pro^_GST, PreScission^pro^, 3CL^pro^ and GST. FT: contain 3CL^pro^, E: elution contains uncleaved 3CL^pro^_GST, PreScission^pro^ and GST. **(C)** SDS 15% Gel of 3CL^pro^ after size exclusion chromatography. **(D)** 3CL^pro^ Chromatogram after size exclusion chromatography. **(E)** CD spectrum of 3CL^pro^.

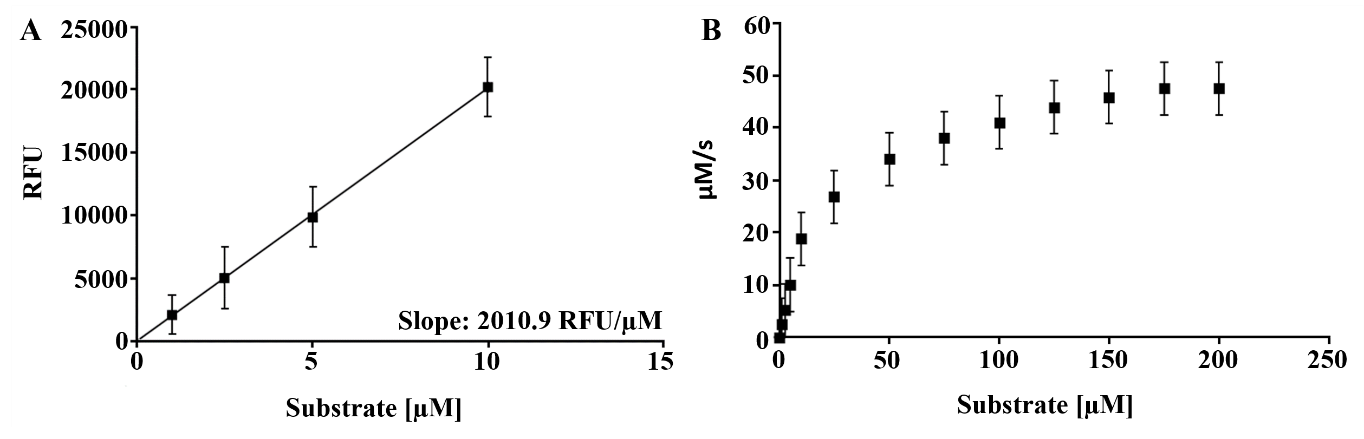

**Figure S2 K_M_ determination of SARS-CoV-2 3CL^pro^. (A)** Standard curve to convert RFU to the amount of the cleaved substrate (µM). **(B)** Michaelis–Menten plot of 0.5 µM protein with various concentrations of FRET substrate. Curve fitting with Michaelis-Menten equation gave the best fitting values of K_M_ and V_max_ as 25.47 ± 3.43 µM and 47.52 ± 2.91 µM/s, respectively. Data represent the mean of triplicate measurements and SD bars are indicated.

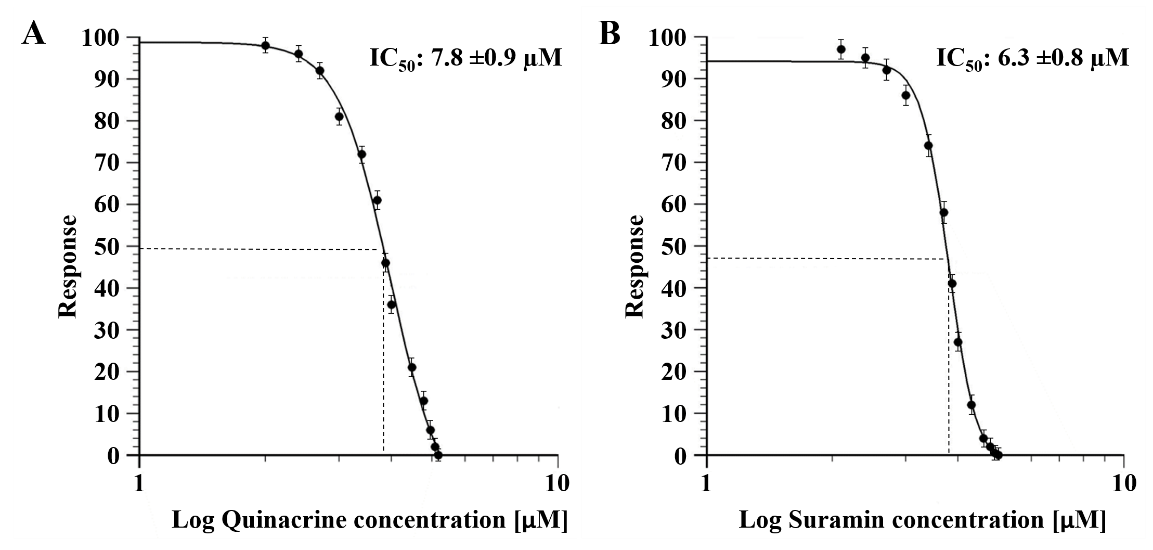

**Figure S3.** **Dose response curves for IC_50_ determination.** The normalized response [%] of SARS-CoV-2 3CL^pro^ is plotted against the Log of the quinacrine and suramin concentration. The determined IC_50_ values are presented within the corresponding picture. **(A)** Dose response curve of quinacrine and SARS-CoV-2 3CL^pro^. **(B)** Dose response curve of suramin and SARS-CoV-2 3CL^pro^. Data represent the mean of triplicate measurements and SD bars are indicated.

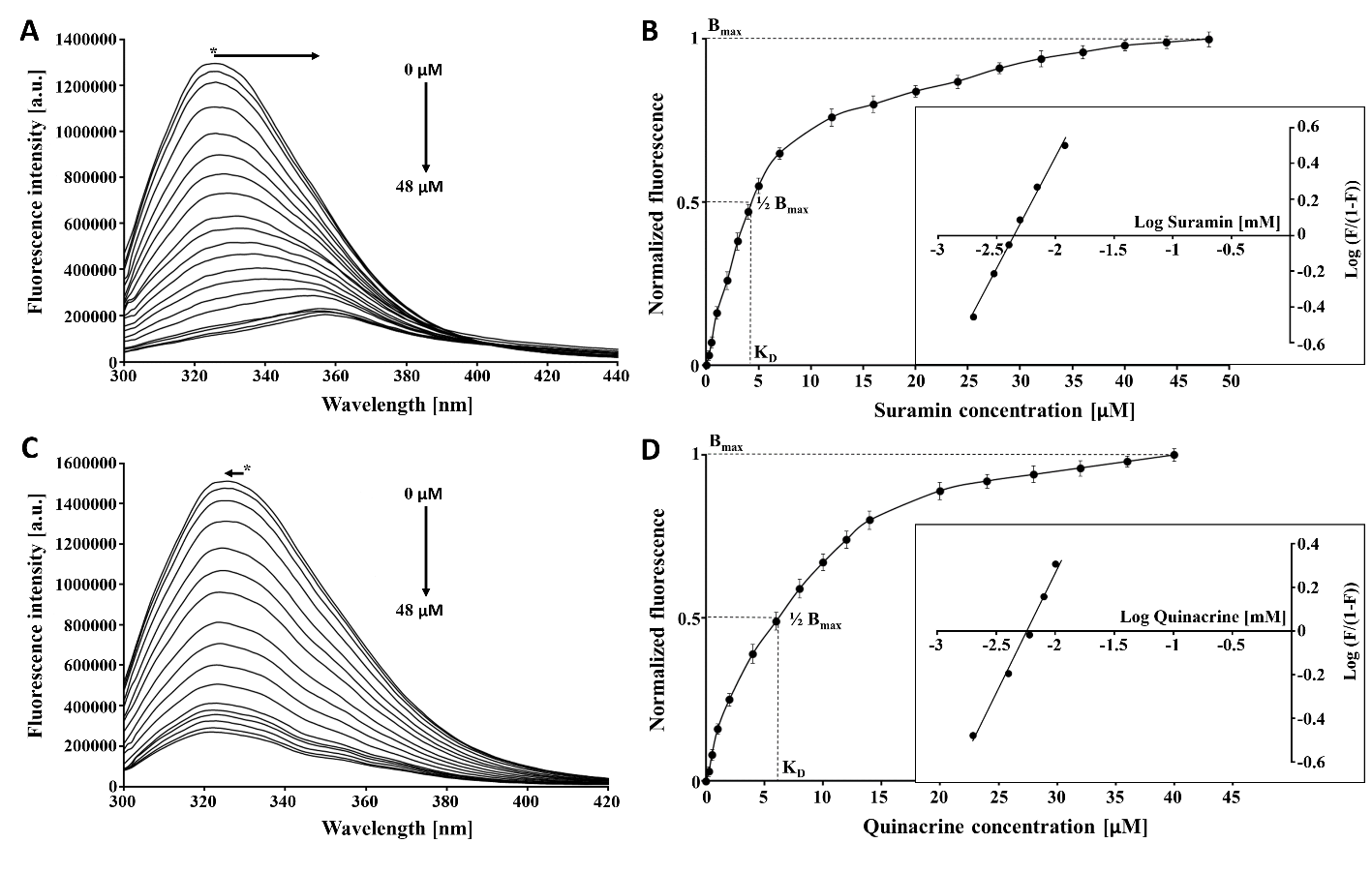

**Figure S4.** **Fluorescence spectroscopy of Trp at 295 nm of SARS-CoV-2 3CL^pro^ in the presence of the suramin and quinacrine. (A)** Fluorescence of SARS-CoV-2 3CL^pro^ under influence of suramin titration demonstrated a red excitation shift of visible Trp (*). **(B)** Binding saturation curve and modified Hill equation determined a K_D_ value of 4.5 ± 1.2 µM for the SARS-CoV-2 3CL^pro^-suramin interaction. **(C)** Fluorescence of SARS-CoV-2 3CL^pro^ under influence of quinacrine titration demonstrated a blue excitation shift of visible Trp (*). **(D)** Binding saturation curve and modified Hill equation determined a K_D_ value of 6.0 ± 1.8 µM for the SARS-CoV-2 3CL^pro^-quinacrine complex. Data represent the mean of triplicate measurements and SD bars are indicated.

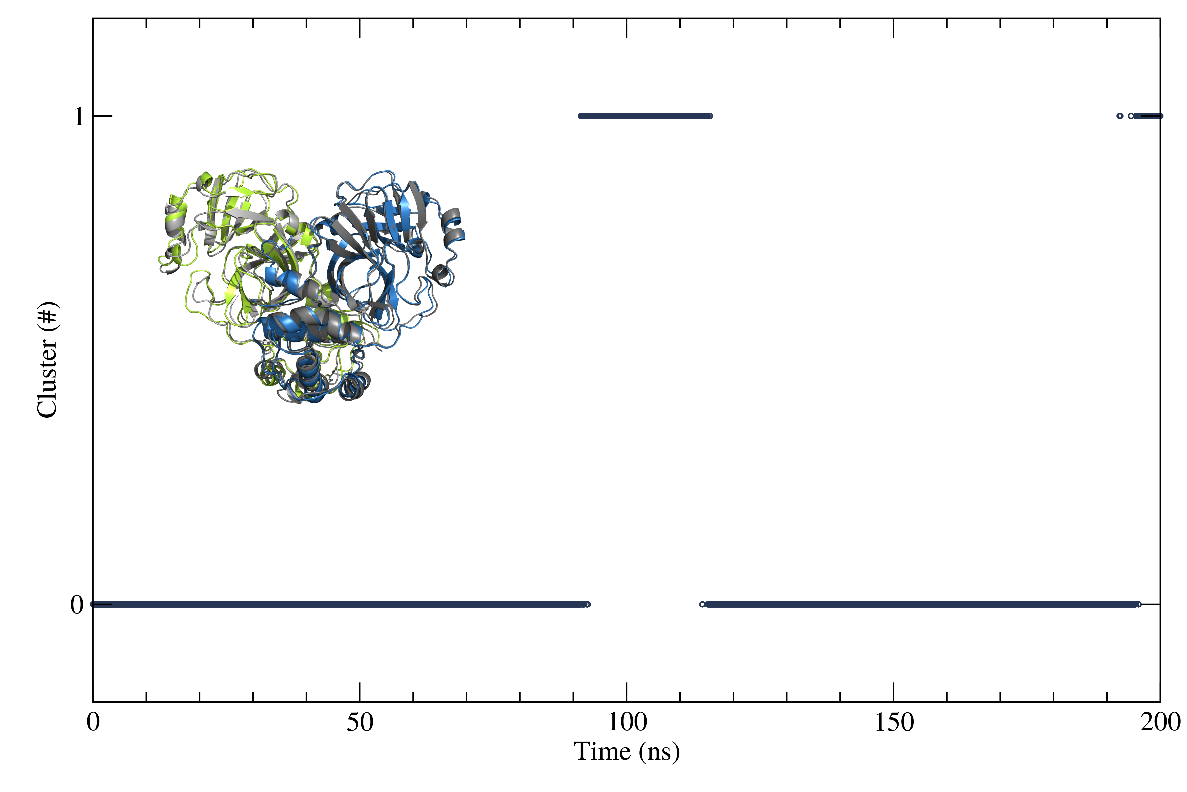

**Figure S5.** **Clustering analysis of the SARS-CoV-2 3CL^pro^ structure after MD simulation.** Pure protein pointed to a chosen structure (cluster 0) that was representative for over 86,1% of the simulation and appeared around 169 ns. The representative structure (C0) is in green and blue, while the (C1) is colored in grey.

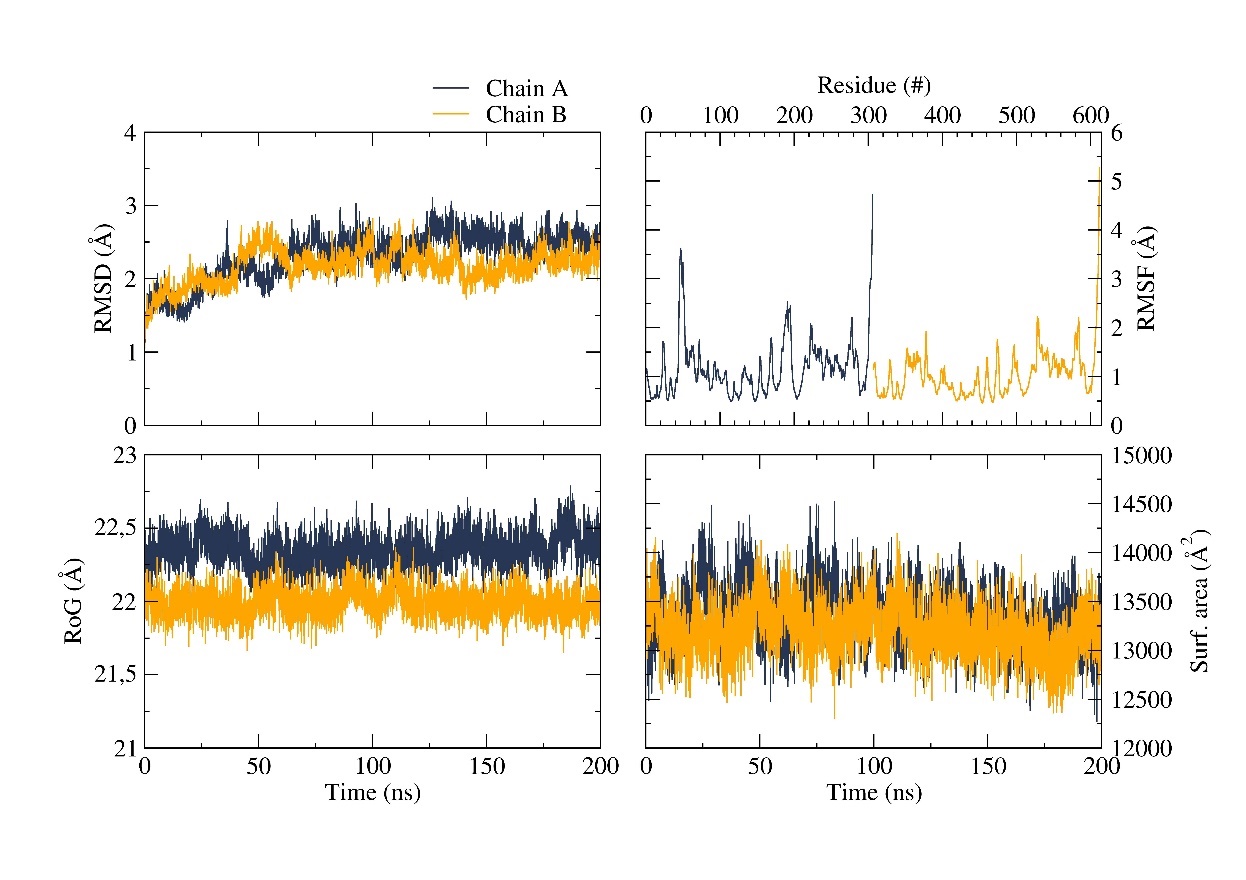

**Figure S6.** **Time dependent modifications of the SARS-CoV-2 3CL^pro^ dimer.** Protomer 1 (blue) and protomer 2 (yellow). RMSD, RoG and surface area as function of time. RMSF for each amino acid.

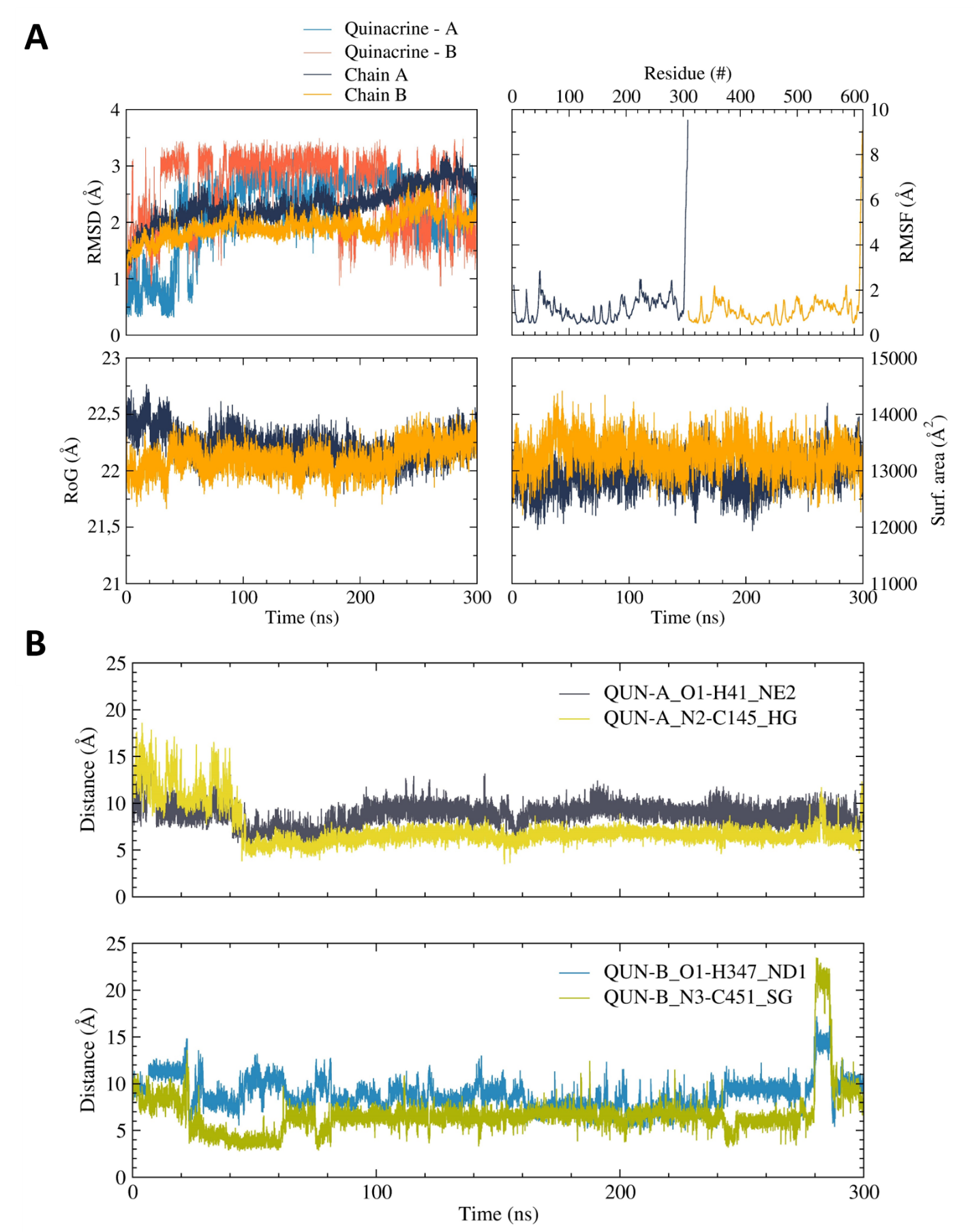

**Figure S7.** **Time dependent modifications of SARS-CoV-2 3CL^pro^ dimer in the presence of quinacrine in each active site.** **(A)** Protomer 1 (blue), quinacrine A (light blue), protomer 2 (yellow) and quinacrine B (pink). RMSD, RoG and surface area as function of time. RMSF for each amino acid. **(B)** Distance over time of quinacrine A and B in the protease active site.

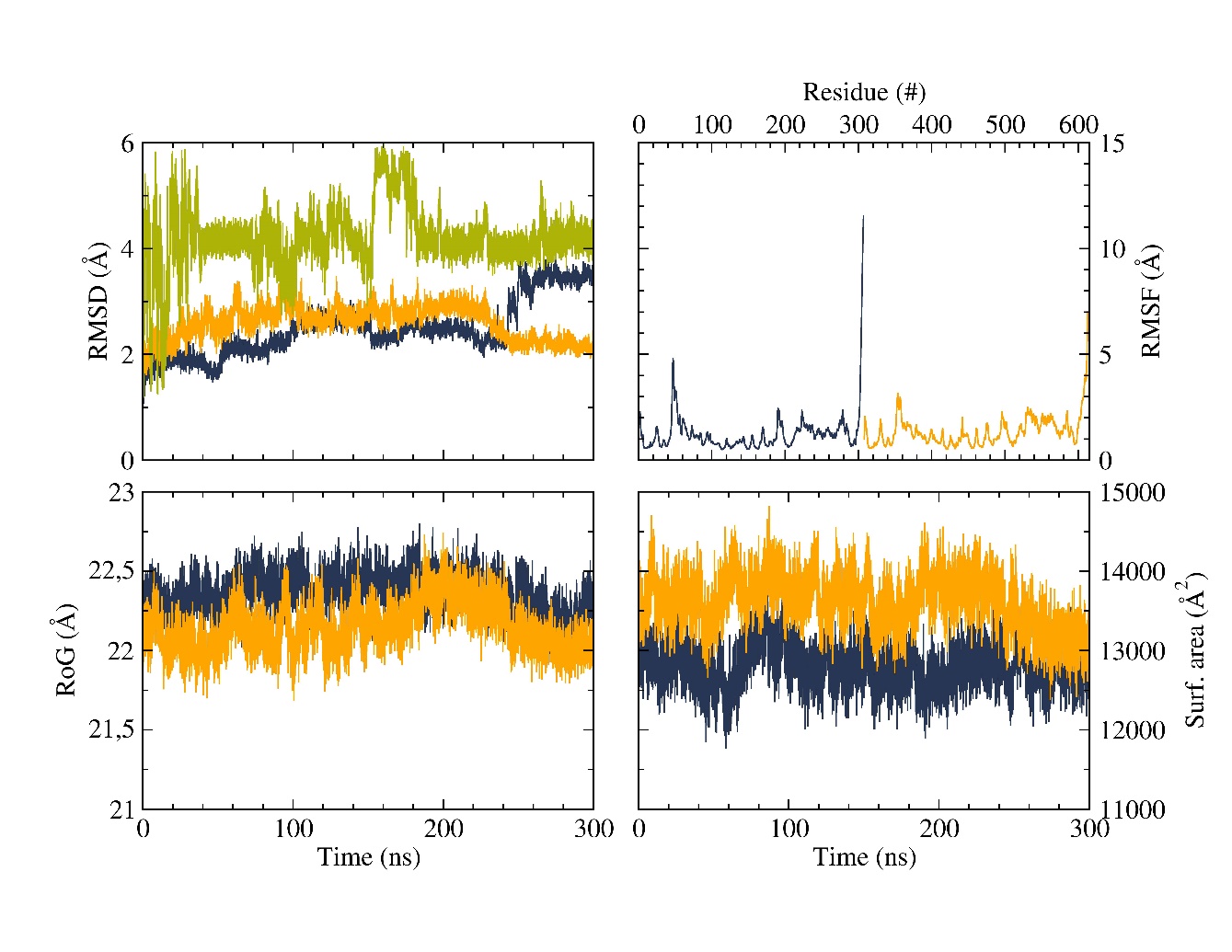

**Figure S8. Time dependent modifications of SARS-CoV-2 3CL^pro^ dimer in the presence of suramin.** Protomer 1 (blue), protomer 2 (yellow) and suramin (green). RMSD, RoG and surface area as function of time. RMSF for each amino acid.

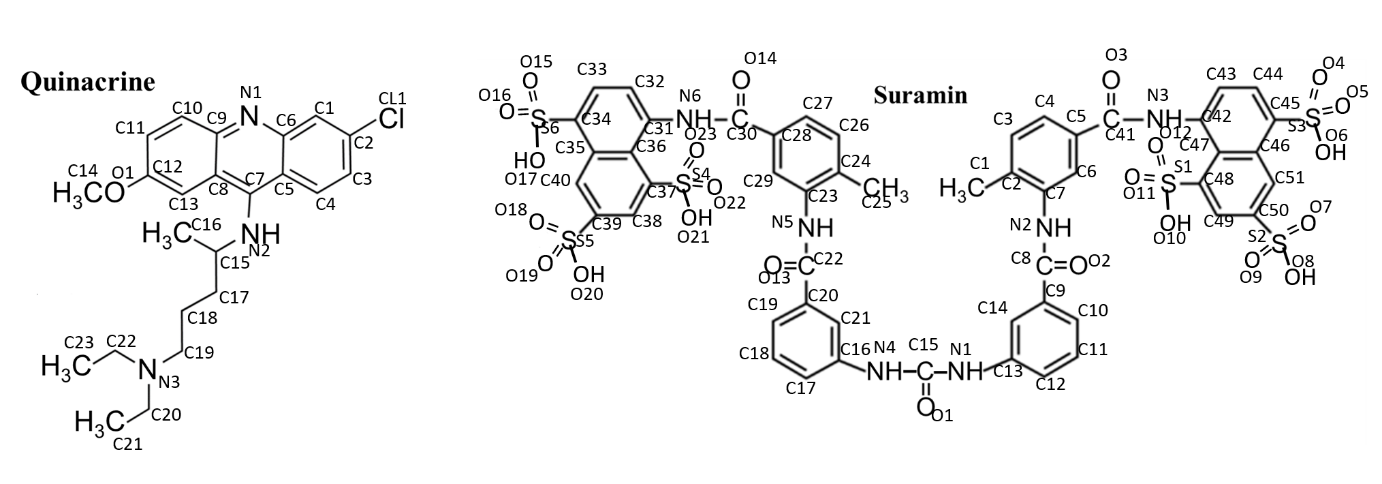

**Figure S9.** **Atom numbers of quinacrine and suramin.**

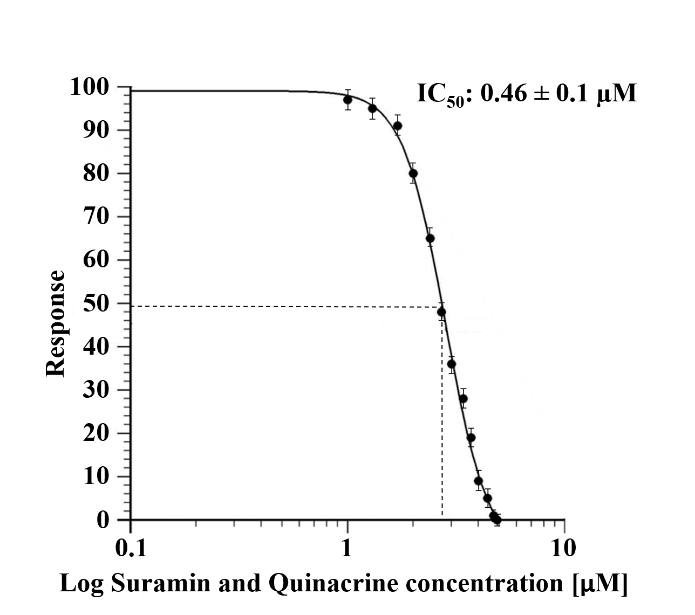

**Figure S10.** **Dose response curve for IC_50_ determination.** The normalized response [%] of SARS-CoV-2 3CL^pro^ is plotted against the Log of the combined quinacrine and suramin concentration. The determined IC_50_ value is presented within the figure. Data represent the mean of triplicate measurements and SD bars are indicated.

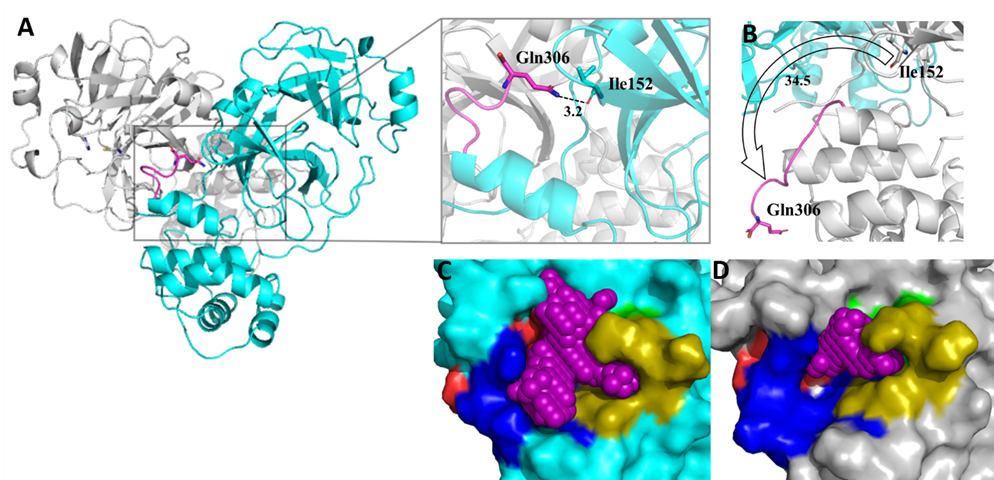

**Figure S11.** **SARS-CoV2 3CL^pro^ dimeric structure after MD simulation.** **(A)** SARS-CoV-2 3CL^pro^ in ribbon view, protomer 1 in turquois and protomer 2 in grey. H-bond between C-terminus Gln306 and Ile152 in protomer 1 is highlighted. **(B)** Protomer 2 Gln306 and Ile152 develop no H-bond. **(C)** Protomer 1 active site pocket volume yields around 261 Å^3^, the substrate binding subsites are highlighted. Green S1`; gold S1; red S2 and blue S3. **(D)** Protomer 2 active site volume yields about 142 Å^3^.

**Table S1.** Contributing amino acids in the

Substrate-binding site of SARS-CoV-2 3CL^pro^.

| Substrate subsite* | SARS-CoV-2  3CL^pro^ |
| --- | --- |
| S1` | His41 |
|  | Gly143 |
|  | Ser144 |
|  | Cys145 |
| S1 | Phe140 |
|  | Leu141 |
|  | Asn142 |
|  | His163 |
|  | Glu166 |
| S2 | His41 |
|  | Met49 |
|  | Tyr54 |
|  | His164 |
|  | Asp187 |
|  | Arg188 |
| S3 | Met165 |
|  | Glu166 |
|  | Leu167 |
|  | Gln189 |
|  | Thr190 |
|  | Gln192 |
| Oxyanion hole | His41 |
|  | Cys145 |

* Based on Tang et al. 2020 [1]
